# Motif-Centered Analyses Reveal Universal and Tissue-Specific Mutagenic Mechanisms Operating in the Human Body

**DOI:** 10.1101/2025.10.07.681015

**Authors:** Safia Mahabub Sauty, Yun-Chung Hsiao, Leszek J. Klimczak, Dmitry A. Gordenin

## Abstract

Somatic mutations are inevitable in human genomes and can lead to tumorigenesis, yet baseline mutagenesis in non-cancerous normal cells remain poorly understood. Here, we analyzed the mutation profiles of 11,949 normal samples across 25 tissues obtained from whole-genome and whole-exome sequencing datasets. We applied stringent statistical hypothesis for detecting enrichment and enrichment-adjusted Minimal Estimate of Mutation Load (MEML) in trinucleotide motifs preferred by known mutagenic processes. We found several cancer-associated mutational motifs in cancer-free tissues. Samples enriched with C→T mutations in nCg motif associated with clock-like spontaneous meCpG deamination were detected across all tissues. We revealed another clock-like motif, T→C substitutions in aTn motif associated with exposure to small epoxides and other S_N_2 electrophiles, in several tissues. Donors with several non-cancerous diseases showed significantly higher, age-independent, and concordant accumulation of aTn and nCg motifs compared to healthy donors. Motifs associated with chemical exposures showed sporadic, tissue and disease-specific mutagenesis. APOBEC-induced C→T and C→G mutations in tCw motif were enriched in bladder, lung, small intestine, liver, and breast with preference for APOBEC3A-like mutagenesis in most. Together, our analyses elucidated several ongoing mutagenic processes in normal human tissues and provided a robust analytical framework for identifying mutagenic sources from somatic mutation catalogues.

## Introduction

Mutations are inevitable in genomes of all species, including humans. Genome sequencing breakthrough more than a decade ago unveiled a massive burden of *de novo* mutations in human cancers (1, 2, 3). Earlier research also led to propose that mutation accumulation in non-cancerous normal somatic cells can be carcinogenic (4, 5). Somatic mutations can also result in disease other than cancer (6, 7, 8). The possible roles of somatic mutagenesis in cancer and other diseases led to questions about the rates and causes of somatic mosaicism and genome instability in humans. Mutation catalogs of somatic cells and tissues can be obtained from conventional whole genome sequencing of cell clones, deep sequencing of micro biopsies, sequencing of multiple single cells from a tissue or by accurate single-molecule sequencing from a non-clonal population of cells derived from a tissue ((9, 10, 11, 12, 13) and references therein). Catalogs of somatic mutations in normal (non-cancerous) human tissues generated in dozens of independent studies were accumulated in the SomaMutDB database (https://vijglab.einsteinmed.edu/SomaMutDB) (14, 15). The collected large volume of mutation data enabled mutational signature extraction to be applied to the entire SomaMutDB dataset, to its’ sub-sets including samples from specific tissues, to other datasets and then to use these signatures for addressing specific hypotheses (see e.g., (16, 17)). Agnostic signature extraction turned to be a powerful and popular tool allowing indication of the presence of known as well as yet unknown mutational processes operating in human cancers (3, 18). However, at the very inception of this method it had been stated that “Parameters [of signature deciphering] to which solutions are sensitive include the number of operative mutational processes, the strength of their exposures, the degree of difference between mutational signatures, the number of analyzed cancer genomes, the number of mutations per cancer genome, and the number of mutation types that are incorporated into the model” (19). These limitations make the determination of the presence of any given signature in an individual sample dependent on other samples of the cohort(s) used for agnostic signature extraction. Moreover, the limitations can be significant in application to signature extraction from the mutational catalogs of normal somatic cells and tissues, where the number of mutations per sample and the sample size of a cohort can be smaller than in cancers. Standard approach with agnostic signatures derived from catalogs of somatic mutations in normal tissues is to align somatic signatures with canonical COSMIC signatures (https://cancer.sanger.ac.uk/signatures/sbs/ (18)) using signature extractor tools and/or calculating cosine similarity (12, 16, 17). This approach allows to highlight presence of some known mutagenic processes but does not assign statistically evaluated P-values to the prevalence of signature(s) in an individual sample.

We previously developed the knowledge-based motif-centered analytical pipeline P-MACD (https://github.com/NIEHS/P-MACD) complimentary to agnostic signature extraction. The pipeline is set to calculate sample-specific enrichment of mutations in trinucleotide motif(s) that have been identified by mechanistic research to be preferred by a single or by a group of related mutagenic factors (see Materials and Methods). The parameters of motif-specific mutagenesis in a sample are just the two odds-ratios, which allows to calculate sample-specific statistics, apply FDR correction, and estimate mutation load caused by a motif-preferring mechanism in each sample of a cohort. Our and others’ prior studies have demonstrated several utilities of motif-centered analyses. Motif-specific calculations of enrichment and mutation load were used in several studies in parallel with signature analyses (20, 21, 22, 23, 24, 25, 26, 27, 28, 29, 30, 31, 32, 33, 34, 35, 36, 37, 38, 39, 40). For example, it allowed to distinguish between APOBEC3A and APOBEC3B mutation load in human cancers (36, 37, 38), detect APOBEC mutagenesis in a subset of lung tumor samples from smokers where agnostic extraction did not indicate APOBEC mutagenesis (39), and identify minor UV-induced base substitution and InDel mutational motifs as corroborating evidence for UV-mutagenesis in human skin (40). Motif-centered analyses have also revealed statistically significant presence of a motif with sequence identical to COSMIC signature SBS16 in all types of human cancers, while the agnostic extraction assigned SBS16 to only two (at most three) cancer types (41). While the peaks of SBS16 are part of SBS5 which is extracted from all human cancers, SBS16 itself is not extracted from most of cancers where SBS5 was detected (41). Altogether, knowledge-based motif-centered analyses proved useful in several cases, where sample-specific quantitation of the impact of a known mutagenic process in individual samples is sought for.

In this work, we applied motif-centered analyses to mutation catalogues of the vast number of somatic samples curated and made available in SomaMutDB (14, 15). We report findings revealed by motif-centered pipeline applied to mutation catalogs of over ten thousand whole-genome and whole-exome sequenced (WGS and WES) samples from a wide range of normal human tissues obtained from healthy and disease-carrying individuals. We found that enrichment with mechanism-associated motifs previously detected in human cancers also exists in normal tissues. We also revealed association of several motifs with tissue type, donor age, and disease status. The analytical framework presented in our study can be applied to any mutation catalogs of individual somatic samples as well to large sample cohorts.

## Materials and methods

### Whole genome and exome datasets

Catalogues of somatic mutations in healthy tissues from whole genome and exome sequencing mapped to GRCh38 reference genome were downloaded as VCF (Variant Call Format) files from SomaMutDB (URL: https://vijglab.einsteinmed.edu/SomaMutDB/download/) (14, 15). A general description of VCF file format can be found in (https://docs.gdc.cancer.gov/Data/File_Formats/VCF_Format/). Mutation calls from embryonic stem cells were not included in analyses. Studies reporting mutation calls in donor groups instead of individual samples were also excluded. We also excluded one study reporting variants in endometrium, as the available VCF files only contained indel calls. Catalogues of mutations in diseased tissues from whole genome and exome sequencing mapped to GRCh38 reference genome were downloaded as VCF files from the FAQ section (question 10) of SomaMutDB (URL: https://vijglab.einsteinmed.edu/SomaMutDB/documentation/). Samples with history of carcinoma and chemotherapy were excluded. For single cell brain samples in SMSDB04 and SMSDB33, VCF files were available for the putative, high confidence somatic mutations called by linked-read analysis (LiRA) pipeline and used to estimate SNV/neuron (42, 43). Only 36 out of the 212 individual neurons with available VCF files originating from these two studies were re-called with SCcaller to detect all mutations and were made available in the SomaMutDB database. We used the VCF files reporting the putative mutation catalogues for all 212 neurons to report mutation spectrum and perform sample-specific motif-centered analyses. Neurons from SMSDB04 that were re-analyzed in SMSDB33 were only reported once with the sample IDs from SMSDB04. Mutation calls generated in studies not included in the SomaMutDB database were collected from the Supplementary materials of the cited papers (44, 45, 46) and from associated dbGaP studies (40, 47). Coordinates of mutation calls obtained from Supplementary materials and mapped to GRCh37 reference genome were converted to GRCh38 coordinates using UCSC LiftOver tool (URL: https://genome.ucsc.edu/cgi-bin/hgLiftOver). Overlapping mutation calls were removed for clones and single cells originating from the same donor in the same tissue biospecimen. Duplicate calls were recorded once and were assigned to samples with the highest variant allele frequency where such information was available. All VCF files were converted to MAF (Mutation Annotation Format) files using the vcf2maf tool (URL: https://github.com/mskcc/vcf2maf) to perform motif-centered analysis. A general description of MAF file format can be found in (https://docs.gdc.cancer.gov/Data/File_Formats/MAF_Format/).

### Detection of COSMIC SBS signatures in the mutation catalogues of normal tissues

Mutation data for all samples within a tissue or disease group were pooled to generate 96-trinucleotide profiles, which were then refitted with published COSMIC SBS signatures (v3.2) using MutationalPatterns R package (48). A second refitting was performed using SBS signatures contributing >5% of mutations in individual tissue or disease group as the reference set. Relative contribution of signatures in different tissues or disease groups was calculated and plotted. Cosine similarity analyses to test the quality of refitting models were performed using cos_sim_matrix() function between observed and reconstructed 96-channel mutation profiles of each tissue and disease group. Signature data from *de novo* signature extraction and decomposition utilized for signature-motif correlation were collected from the supplementary materials of previous studies (17, 40, 45, 49, 50). Sample specific signature refitting was performed using both MutationalPatterns (48) and SigProfilerAssignment (51) tools. A second refitting was performed with each tool using SBS signatures contributing >5% of mutations in an individual sample as the reference set.

### Detection of enrichment of mutations in trinucleotide motifs and calculation of minimum estimate of mutation load associated with known mutagenic processes

Trinucleotide motifs and their sub-motifs are constructed based on mechanistic knowledge and experimental validation of the mutagenic preferences of the associated mutational processes. Motif-centered analyses to find statistically significant enrichment of specific mutagenic processes were performed using the P-MACD software (URL: https://github.com/NIEHS/P-MACD) as previously described (40, 41, 47, 52). Briefly, enrichment of a mutational motif was calculated by normalizing the incidence of a specific base substitution in a trinucleotide motif to the random incidence of that mutation in the background genomic context. For this calculation, context is defined as ±20 bases surrounding the mutated base to concentrate on the sequenced part of the genome. This allows to capture only localized effects of lesion formation, TLS, and repair, to avoid epigenomic impacts on mutagenesis in the larger genomic regions within each sample, and to circumvent any regional amplification biases introduced by different sequencing platforms. Reverse complements were included in calculations and motifs were reported as pyrimidine nucleotide of a base pair. The mutated base is shown in capital, and IUPAC codes are used for ambiguous nucleotides. The formula to calculate enrichment is shown below using aTn→aCn mutational motif as an example:

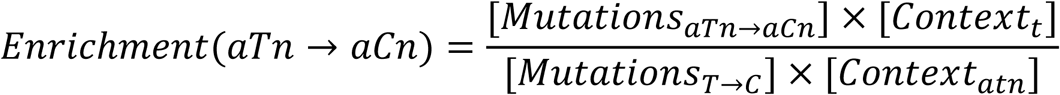

We excluded ‘complex’ mutations, defined as mutations ≤10bp apart, from this calculation as they may arise from the activities of the translesion polymerases and may confound the detection of mutagenic mechanisms with specific motif preferences. We performed one-sided Fisher’s Exact test to statistically evaluate the overrepresentation of mutations in a motif over random mutagenesis in individual samples. Ratio of the number of mutations in motif (*Mutations_(aTn→aCn)_*) to mutations that are not in motif ((*Mutations_(T→C)_*) - (*Mutations_(aTn→aCn)_*)) were compared to the ratio of the number of bases in the context that conformed to the motif (*Context_(atn)_*) to bases that did not conform to the motif ((*Context_(t)_*)-(*Context_(atn)_*)). For samples that show statistically significant enrichment after correction for multiple hypothesis using Benjamini-Hochberg method (enrichment >1, q value ≤0.05), we calculated the minimum estimate of mutation load (referred as MEML) attributable to the mutagenic process associated with the trinucleotide motif. MEML was calculated using the following formula:

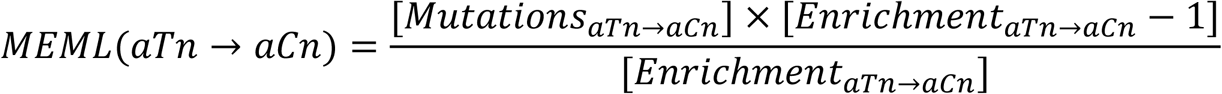

MEML was recorded as zero for samples that did not show statistically significant enrichment of a mutational motif. All figures report sample-specific MEML values unless otherwise indicated.

### Statistical analyses and data visualization

All statistical analyses were performed in R statistical software (version 4.4.0) via RStudio. Spearman correlation was performed for all correlation analyses using cor.test() function. Age correlations were performed with average MEML count of all samples originating from a donor. wilcox.test() function was used to perform all tests of significance, followed by correction for multiple hypothesis by Benjamini-Hochberg method using p.adjust() function. Donors in the dataset were divided in three age terciles, and tests of significance were performed for each age group within individual studies that reported both healthy and diseased cohorts to avoid batch effect and minimize the effect of aging in detecting true difference between healthy and diseased tissues. For single cell brain samples in SMSDB04 and SMSDB33, MEML counts calculated by motif-centered analyses of the putative SNVs were prorated by sample-specific multipliers to estimate genome-wide MEML load. Multipliers were calculated as

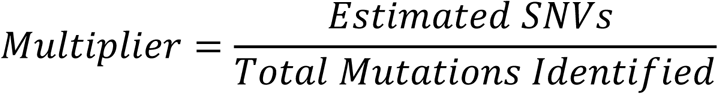

For SMSDB04, numerator values were obtained from the ‘*Estimated SNVs (per autosomes)*’ column and the denominator values were obtained from the ‘*Total Mutations Identified*’ column of Supplementary_table_S5 (42). For SMSDB33, we used values from Supplementary_table_S6 (43). For numerator of MDA-amplified available VCF files, we used values from column ‘*Estimated SNVs (per autosomal genome, post-filtering)*’ of sheet ‘*Rates per Neuron (MDA)*’. For numerator of PTA-amplified available VCF files, we used values from column ‘*Estimated SNVs (per autosomal genome)*’ of sheet ‘*Rates per Neuron (PTA)*’. Values from columns ‘*Phaseable Mutations Identified*’ were used as denominators for all samples from SMSDB33.

Multivariable regression analyses were performed using lm() function using age and disease status as predictor variables and motif-specific MEML as the dependent variable. Unless otherwise indicated, all data visualizations were performed using ggplot2 R package (53) and Adobe Illustrator (54).

## Results

### Study design

We analyzed the mutation calls from WGS and WES samples of non-cancerous (i.e., normal) human tissues. The mutation calls were retrieved from the somatic mutation database SomaMutDB (https://vijglab.einsteinmed.edu/SomaMutDB) (14, 15), and from the supplementary materials of several papers not included into SomaMutDB (see Materials and Methods) (Supplementary Table S1a). The dataset of mutation calls was categorized into two groups, healthy and diseased normal tissues. The mutation calls in the healthy cohort are from 10,625 samples of 777 individual donors, spread across 25 organs (Supplementary Table S1b). 8,054 of these samples are WGS with total 7,623,352 mutations (sample mutation loads range from zero to 34,629). 2,571 healthy normal samples are WES with total 179,208 mutations (sample mutation loads range from zero to 5,957). Detailed information of both WGS and WES healthy samples, including their mutation loads, are presented in Supplementary Table S1b. The mutation calls in the diseased cohort are from 1,323 samples of 223 Individual donors, spread across 6 tissue types and 15 different disease conditions (Supplementary Table S1c). 1,226 of the diseased samples are WGS (with 3,243,116 total mutations; sample mutation loads range from zero to 18,243) and 96 samples are WES (with total 65,843 mutations; sample mutation loads range from 9 to 2,931). Detailed information of the diseased cohort is available in Supplementary Table S1c.

We first compared the mutation profiles of different normal tissues with the published COSMIC reference SBS signatures derived from large cohorts of cancer mutation catalogs. Then, we performed motif-centered analyses to find enrichment of known mutagenic processes and quantify their presence in individual samples. To conduct motif-centered analyses of the mutation calls, we curated a list of trinucleotide motifs preferred by known mutagenic mechanisms and supported by experimental evidence and mechanistic knowledge (Table 1). In order to resolve potential overlaps between motifs centered around the same base substitution, we built a non-overlapping sub-motif, where needed (Supplementary Table S2).

**Table 1:** List of knowledge-derived mutational motifs.

| Motif name<sup>1</sup> | Reference trinucleotide | Mutant trinucleotide | Motif for SBS correlation | Mechanism | References | Corresponding COSMIC signature(s) |
| --- | --- | --- | --- | --- | --- | --- |
| aTn | aTn | aCn | aTn→aCn | Epoxide, S<sub>N</sub>2 electrophiles | (41) | SBS16 |
| yCn | yCn | yTn | yCn→yTn | UV | (40, 47, 55) | SBS7a, SBS7b |
| nCg | nCg | nTg | nCg→nTg | Spontaneous deamination of meCpG | (40, 47, 56) | SBS1 |
| nTt | nTt | nCt | nTt→nCt | UV (minor) | (40, 47) | SBS7c, SBS7d |
| tCw | tCw | tTw | tCw→tTw | APOBEC | (52) | SBS2 |
| tCw | tCw | tGw | tCw→tGw | APOBEC | (52) | SBS13 |
| tgC | tgC | tgA | tgC→tgA | Redox | (57)) | SBS18 |
| hTg | hTg | hGg | hTg→hGg | S<sub>N</sub>2 alkylation | (58) | NA |
| aCy | aTy | aCy | aCy→aTy | S<sub>N</sub>1 alkylation | (58) | SBS11 |
| gCn | gCn | gAn | gCn→gAn | Acetaldehyde | (59) | NA |
| cTg | cTg | cAg | cTg→cAg | Aristolactam (from Aristolochic acid) | (60) | SBS22a, SBS22b |

1. Bases within each motif are shown in IUPAC codes

We evaluated statistically significant over-representation of 11 knowledge-based trinucleotide motifs using motif-centered analysis tool P-MACD (https://github.com/NIEHS/P-MACD). P-MACD calculates enrichment of mutations in the trinucleotide context relative to random mutagenesis. It assigns sample-specific P-values and calculates enrichment-adjusted Minimum Estimate of Mutation Load (hereafter referred to as MEML) for samples that have a statistically significant enrichment of mutational motif after correction for multiple-hypothesis testing. By forming a stringent statistical hypothesis and performing sample-specific analyses, we extracted the MEML value that can be attributed to a specific mutagenic mechanism. Conveniently, non-zero MEML value indicates statistically significant enrichment with a motif, so these two terms can be used interchangeably.

### Mutation spectrum and the contribution of COSMIC SBS signatures in normal tissues

Dataset of mutations in healthy normal tissues contained a total of 7.8 million single base substitutions (SBS) identified by whole exome or by whole-genome sequencing (hereafter referred as WES or WGS, respectively). Median SBS counts in WGS healthy tissues ranged from 22 to 4,025, while counts in WES healthy tissues ranged from 9 to 499 (Fig. 1A, Supplementary Table S3a). Similar to prior reports of genome-wide mutagenesis in multiple cancer types (1, 3, 18) we observed highest fraction of C to T substitutions across all tissues. Mutation catalogs of normal tissues with noncancer diseases contained 3.3 million substitutions with median SBS count ranging 29 to 6,026 in WGS samples and 35 to 1,135 in WES samples (Fig. S1A, Supplementary Table S3b).

**Figure 1:**
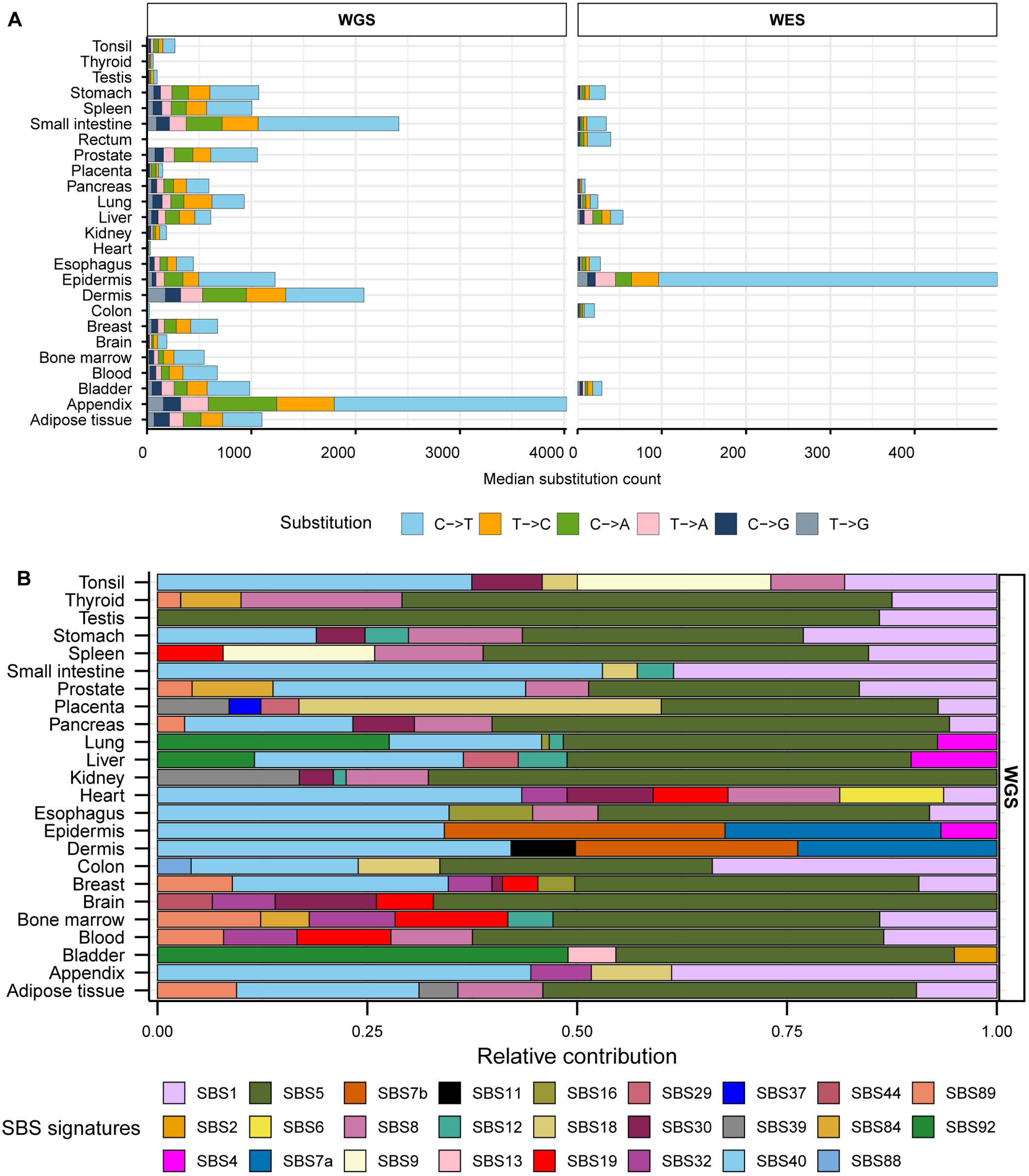
Somatic mutation spectra and contribution of COSMIC signatures in healthy tissues. **A**. Median counts for each base substitution including reverse complements are shown in descending order for healthy WGS and WES samples. **B.** Relative contributions of published COSMIC signatures in the mutation profiles of WGS healthy tissues. All source data are available in Supplementary Table S3.

We first performed mutation signature analysis to understand the patterns of mutagenesis in the healthy and diseased tissue cohorts. We limited signature analysis to refitting with published COSMIC SBS signatures and did not perform *de novo* signature extraction due to the extraction method’s limitations outlined in the Introduction. We generated 96-channel mutation profiles using all mutation calls from each healthy and disease tissue types and fitted them with COSMIC SBS signatures derived from PCAWG cancer mutation catalogues (18) to estimate the relative contribution of cancer-associated signatures in the mutation loads of normal tissues (Supplementary tables S3c, S3d). In consensus with the originating studies that performed *de novo* signature extractions (17, 45, 49, 50, 61, 62), we detected clock-like signature SBS5 with no assigned etiology to be the largest contributor in most healthy (18 out of 24) and diseased (10 out of 14) WGS tissue types (Fig 1B, Supplementary Fig. S1,B). SBS5 was also detected in two WES healthy tissues ( Supplementary Fig. S2A). Another flat signature with no known etiology, SBS40, was readily detected in multiple healthy and diseased WGS tissue types (Fig 1B, Supplementary Fig. S1B). Combined contribution of these two signatures ranged from 0% to 86% of the total mutation pool of different healthy and diseased tissues (Supplementary table S3e).

We detected the contribution of SBS1 associated with meCpG deamination in 17 out of 24 WGS healthy tissues and 9 out of 14 WGS disease types (Fig 1B, Supplementary Fig. S1B, Supplementary tables S3c, S3d). Analyses of WES samples revealed SBS1 in 5 out of 10 healthy and 1 out of 2 disease tissues (Supplementary Fig S2A). UV-associated signatures SBS7a and SBS7b were detected in epidermis and dermis WGS and WES tissues (Fig 1B, Supplementary Fig. S1B, S2A). SBS7b was also detected in stomach, rectum, esophagus, and colon WES tissues where UV-mutagenesis is not expected (Supplementary Fig. S2A). SBS2 and SBS13, signatures associated with deamination by APOBEC enzymes, were detected to have small contributions only in healthy bladder tissue. We assessed the quality of refitting model by performing cosine similarity analyses between the observed 96-trinucleotide mutation profiles and the profiles reconstructed from estimated contributions of COSMIC signatures. We detected cosine similarity value >0.9 for all healthy and diseases tissue groups across both WGS and WES samples, suggesting reliable reconstruction by the refitting models (Supplementary table S3f).

We noted that the spontaneous deamination of meCpG dinucleotides, which is a known endogenous mutator in human cells, was not reliably identified in all tissues analyzed by signature refitting. In contrast, signatures associated with tissue-specific mutagenesis were reliably detected only in tissues with a high likelihood of exposure, like UV-mutagenesis in skin. Additionally, analyses of WES samples with low number of mutations resulted in assignment of signatures to biologically implausible tissue types. Hence, we proposed that our framework of motif-centered analysis may validate existing mutagenic sources, reveal undetected mechanisms, and quantify sample-specific contributions of known mutagenic sources where signature analysis may be underpowered or unsuitable (see Introduction).

### Mutational motifs derived from mechanistic research are enriched in various human tissues and correlate with mutation signatures

We used P-MACD pipeline to assess enrichment of eleven selected trinucleotide mutational motifs in the mutation catalogues of all individual samples. These mutational motifs were previously associated with mutagenic mechanisms indicated in mechanistic studies and were also found in human cancers and in normal human skin (Table 1). Strong experimental support is evident from previous studies for nCg motif characteristic of cytidine deamination in meCpG dinucleotides (56), tCw motifs of APOBEC mutagenesis (52, 63, 64), aTn motif of exposure to small epoxides (41), and yCn motif of UV-mutagenesis (40, 47, 55). We analyzed all WGS samples, as well as WES samples which have 100-fold less mutations compared to the WGS samples.

Analysis of mutational motif enrichment indicated wide-spread presence of the two motifs associated with mutagenic processes in different types of morphologically normal healthy tissues (Fig. 2A). The aTn and nCg, were detected in 4%-100% and 43%-100% of the healthy WGS samples, respectively (Supplementary Table S4a). Statistically significant enrichment with nCg was also detected in 9%-93% of WES samples, while 0%-20% WES samples showed aTn MEML, further validating the robustness of the motif-centered analysis (Supplementary Table S4a, Supplementary Fig. S2B). aTn and nCg were also detected across all diseased tissues (Fig. 2B). aTn was 3-100% of the WGS samples and 4%-12% of the WES samples. nCg MEML was detected in 26%-100% of WGS and 94%-100% of WES diseased samples (Fig. 2B, Supplementary Fig. S2C, Supplementary Table S4b). Other motifs were detected only in some organs and diseased tissues, suggesting tissue-specificity of the underlying mutagenic processes. UV-associated motifs yCn and nTt were detected in multiple tissues with the largest contributions in dermis and epidermis (Supplementary Table S4a, b). C→T and C→G mutations in APOBEC-associated motif tCw were also detected in multiple tissue and disease types (Supplementary Table S4a, b).

**Figure 2:**
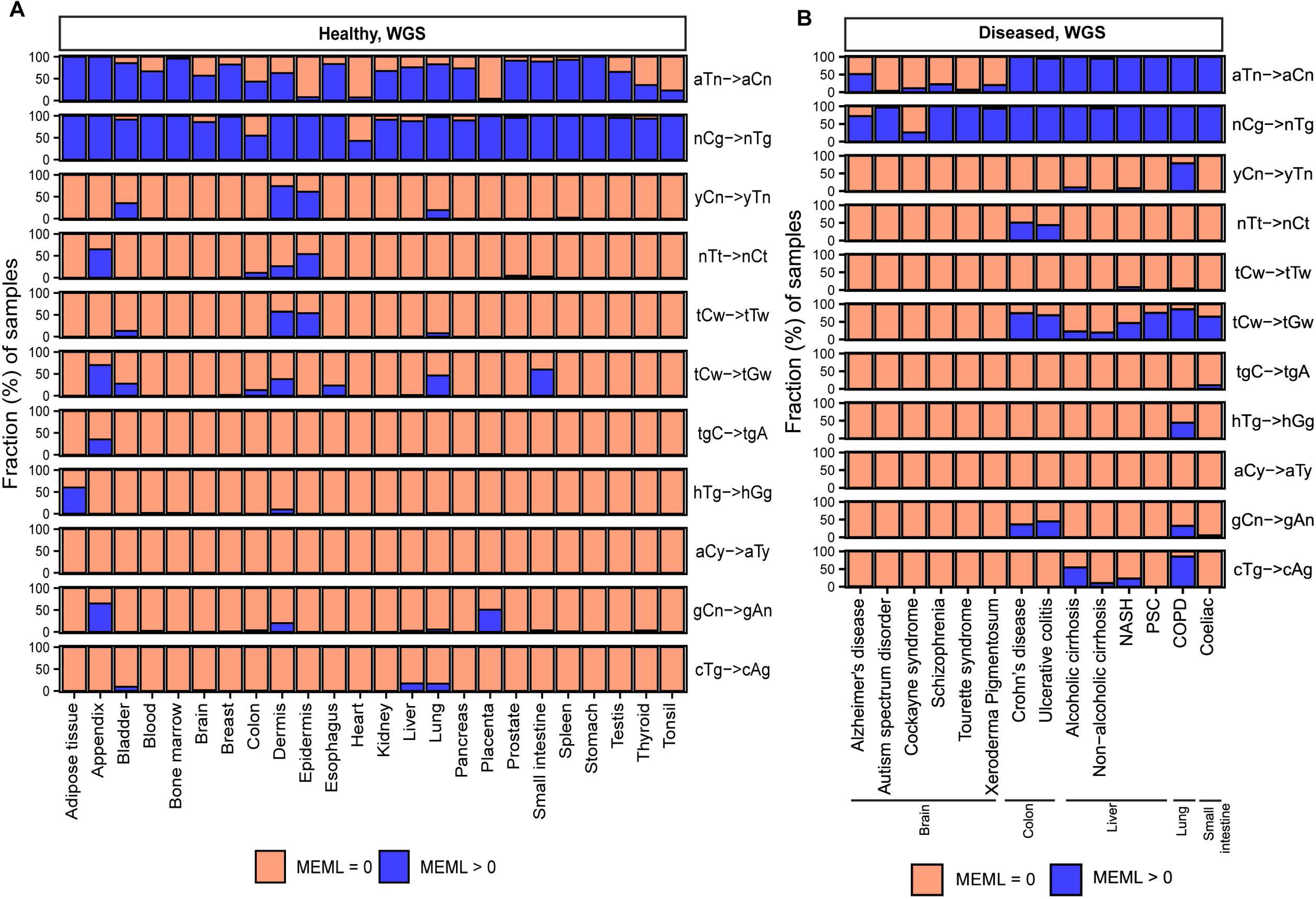
Prevalence of knowledge-based motifs in normal tissues. The percentages of samples with zero and non-zero MEML counts for the 11 known mutational motifs are shown for (**A**) healthy and (**B**) diseased tissues. Only WGS samples are shown. WES samples are shown in Supplementary Fig. S3. All source data are available in Supplementary Table S4a, b.

To understand the inter-method agreement between the motif-centered estimates of mutagenesis in mechanistically-linked mutational motifs and mutation load assigned by agnostic extraction of mutation signatures, we sought to perform Spearman correlation analyses between motif MEML and signature exposures for 1,386 healthy samples (1,310 WGS, 76 WES) where sample-specific signature data were reported in the originating studies (17, 40, 45, 49, 50). We performed correlation analyses across all 11 knowledge-derived mutational motifs and all SBS signatures available within each tissue category (Supplementary table S4c). We found that the two ubiquitous motifs aTn and nCg showed significant positive correlation with signatures SBS1 and SBS5 in most tissues analyzed (Fig. 3A, B). Both SBS1 and SBS5 are clock-like signatures (18). SBS1 signature is analogous and mechanistically concordant to the clock-like motif nCg→nTg, whereas SBS5 contains peaks for all 96 trinucleotide motifs including aTn and nCg motifs (Supplementary Fig. S3A). aTn motif has previously been shown to be clock-like in cancers and exhibited significant positive correlations with both SBS1 and SBS5 in the PCAWG cancer cohort (41). The strong correlation of aTn and nCg motifs with both SBS1 (close match to nCg motif) and SBS5 (flat signature) observed in normal cells is likely driven by their shared clock-like properties, as well as partial or total sequence similarity between these motifs and signatures (Fig. 3A, B, Supplementary Fig. S3A). Interestingly, SBS16 signature with close match to aTn motif (Supplementary Fig. S3A) was not detected by agnostic extraction in these samples.

**Figure 3:**
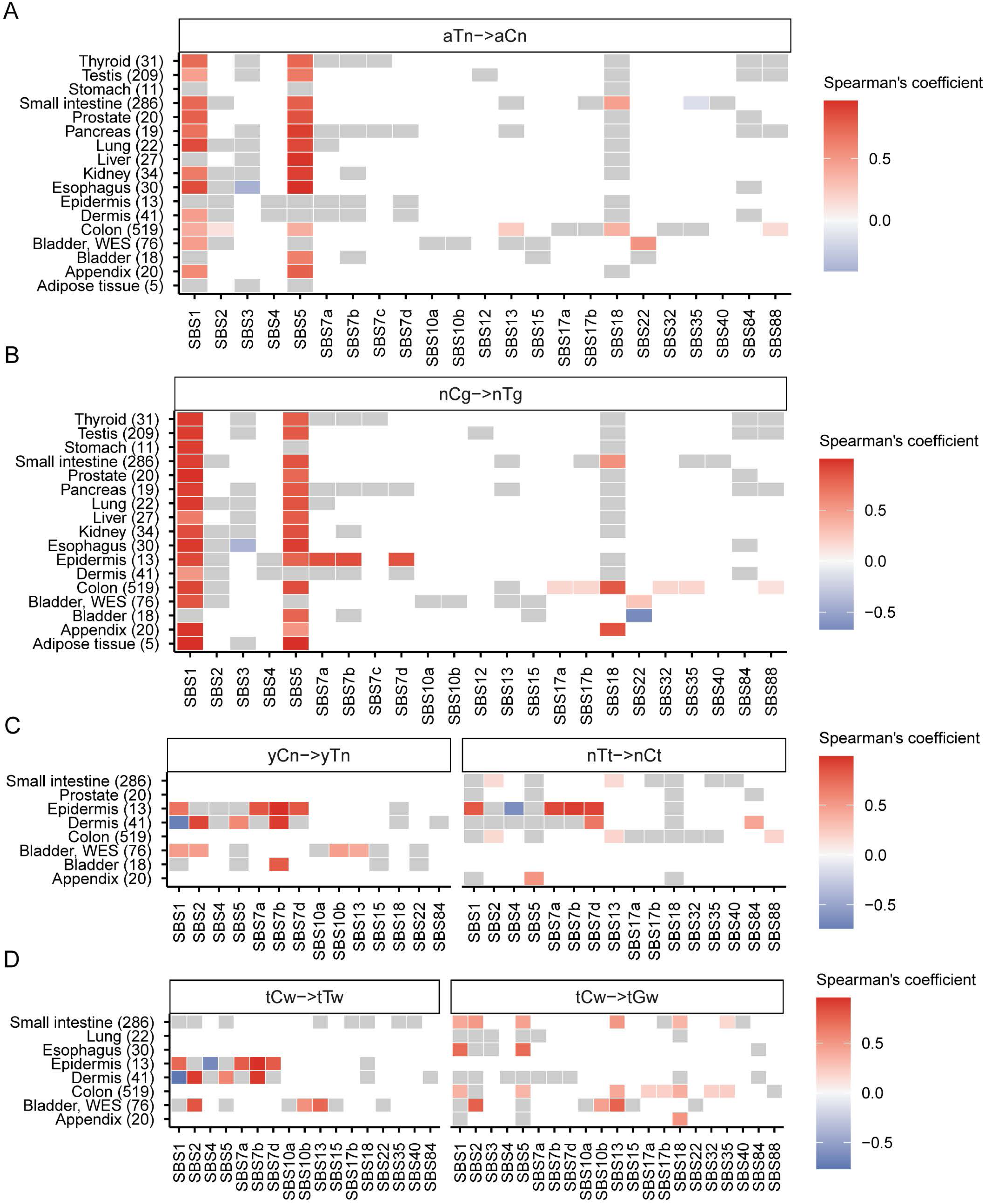
Correlation between mutational motifs and signatures. Correlation matrix showing Spearman’s *ρ* for agnostically extracted COSMIC SBS signatures vs mutational motifs associated with **(A)** exposure to small epoxides and S_N_2 electrophiles (aTn→aCn), **(B)** spontaneous deamination of meCpG (nCg→nTg), **(C)** UV-mutagenesis (yCn→yTn and nTt→nCt), and **(D)** APOBEC-mutagenesis (tCw→tTw and tCw→tGw) for healthy tissues indicated on y-axis. Numbers in parenthesis indicate total number of samples in different tissue types. Grey boxes indicate statistically insignificant correlations after correction for multiple hypothesis using Benjamini-Hochberg method. All source data are available in Supplementary Table S4c. Profiles of COSMIC signatures used for correlations with motifs are shown on Figure S3.

We also detected significant correlations of UV-associated motifs yCn and nTt with UV-associated signatures SBS7a, b, d in dermis and epidermis tissues (Fig. 3C). yCn motif also showed spurious correlations with SBS1 and SBS5 in some tissues, likely due to its complete sequence overlap with nCg motif (Supplementary Fig. S3B). Spurious correlations of nTt motif with SBS1 and SBS5 may be indicative of non-UV lesion motifs that overlap with aTn (Fig. 3C). We detected significant positive correlation between yCn motif and APOBEC-associated signature SBS2 in bladder as well as in dermis (Fig. 3C). This may stem from the overlapping motif preference of UV-induced lesions and APOBEC-mediated deamination of cytosines (see later sections) (Supplementary Fig. S3B). In the same vein, we detected significant correlation between APOBEC-associated motif tCw→tTw with UV-signatures in epidermis (SBS7a, SBS7b, SBS7d) and dermis (SBS7a) (Fig. 3D). We detected significant correlation of both APOBEC-associated motifs with both APOBEC-associated signatures only in WES bladder tissue (Fig. 3D). APOBEC-related tCw→tGw motif aligning with SBS13 showed strong correlations with SBS13 in all tissues where both the motif and the signature were detected. It also correlated with SBS2 reflecting the T→C branch of APOBEC mutagenesis in bladder and in small intestine (Fig. 3D, Supplementary Fig. S3B).

To evaluate the sensitivity of detection of mutagenic activities in individual samples, we performed signature refitting within the mutation catalogues of individual samples using two refitting tools, MutationalPatterns (MP) (48) and SigProfilerAssignment (SPA) (51). We plotted the percentage of samples in both healthy and diseased cohorts in which SBS1 and SBS16 were detected by the signature refitting tools, along with the percentage of samples in which their corresponding motifs (nCg and aTn, respectively) were detected using P-MACD, the tool for motif-centered analysis (Supplementary Fig S4). Percentage of samples with detectable contribution of other signatures with corresponding motifs are available in Supplementary Table S4d.

We observed algorithm-dependent variability in the estimated contributions of both SBS1 and SBS16. SPA consistently detected SBS1 in a higher proportion of samples across most tissues in both healthy and diseased cohorts, whereas MP showed greater sensitivity for SBS16 (Supplementary Fig. S4). Detection of nCg motif by P-MACD was broadly comparable to SBS1 detection by SPA; however, motif-centered analyses identified aTn motif in more samples than SBS16 was detected by either refitting tool. We also identified instances of signature misassignment at the individual-sample level that were not apparent in cohort-level analyses. For example, UV-associated signatures SBS7a and SBS7b were detected in samples across multiple tissues beyond dermis and epidermis (Supplementary Table S4d), where they were exclusively observed in the cohort-level analysis (Fig. 1). This discrepancy likely arises because individual samples have fewer mutations, making the observed mutation patterns more susceptible to random variation in mutation counts and types than in the cohort-level analyses. While P-MACD also detected UV motifs in biologically implausible tissues (Fig 2), such misassignments can be resolved using sub-motif analyses and motif-motif correlation (see later section).

Together, these findings showed that while signature assignment by refitting to pre-defined signatures can be performed at single sample level, the results would be highly tool-dependent which would not allow reliable cross-cohort statistical comparisons. Motif analyses can complement signature analyses to cross-validate biological interpretations where signature etiology is understood and offer mechanistic knowledge for signatures with yet unknown etiology (e.g., SBS16). Below we dissect the features of individual motifs in normal healthy tissues and highlight the utility of motif-centered analyses to disentangle confounding mutagenic mechanisms. We then explore association of motif-specific mutagenesis with disease state of noncancer tissues.

### The small epoxide mutagenesis motif aTn→aCn is enriched in samples across all human tissues and accumulates along the lifetime

The aTn→aCn motif (referred to aTn throughout this paper) was experimentally identified as a defining feature of hypermutation caused in a yeast sub-telomeric single-stranded DNA (ssDNA) model system in response to treatment with small epoxide glycidamide – an electrophile reacting with nucleobases via S_N_2 mechanism (41). Glycidamide-induced hypermutation in ssDNA showed clear overrepresentation of A→G substitution in nAt context (T→C in aTn in reverse complement). The base preference of glycidamide showed similarity with the chemical preference of S_N_2 reacting electrophiles in ssDNA and with the mutagenic preference of another electrophilic alkylating agent reacting via S_N_2 mechanism, methyl methanesulfonate (MMS), supporting the hypothesis about an aTn motif preference not only by small epoxides, but by a broader class of S_N_2 reacting electrophiles ((41, 58) and references therein). Analysis of mutation catalogs from PanCancer Analysis of Whole Genomes (PCAWG) (32) detected widespread enrichment and clock-like accumulation of aTn motif in multiple cancer types. Ubiquitous nature of aTn motif further suggested that it may be a feature of a broader class of mutagens instead of glycidamide alone. Prevalence of aTn motif was shown to have strong positive correlations with mutation loads assigned to COSMIC signatures SBS1, SBS4, SBS5 and SBS16 in tumors where these signatures were detected. In cancers with smoking as a risk factor, aTn mutation load showed a strong and significant association with the smoking history of the patients. Smoking-related aTn mutation load in lung tumors largely exceeded age-dependent presumably endogenous level of mutagenesis. Age-dependent aTn accumulation was also detected in healthy skin and brain cells (41). Given its newly identified ubiquity in normal tissues (Fig 2), we systematically dissected the features of aTn motif to comprehensively characterize the activity of this mutagenic process in normal human tissues.

We detected aTn motif prevalence in morphologically healthy samples across all tissue types. While aTn→aCn MEML contributed only a modest fraction of total AT pair mutations (A→C, A→G, A→T, including reverse complements), its’ statistically significant enrichment was detected in WGS samples across all tissue types (Fig. 4A, Supplementary Table S5a). This indicates the motif’s utility for evaluating the level of epoxide- or even broader class of S_N_2 electrophile-induced mutagenic exposure. The aTn motif-associated mutagenesis was also detected in WES samples, albeit at lower prevalence as compared to WGS mutation catalogs. Multiple samples showing non-zero aTn MEML were detected in all tissue types except epidermis and heart, (Fig. 4B, Supplementary Table S5b).

**Figure 4:**
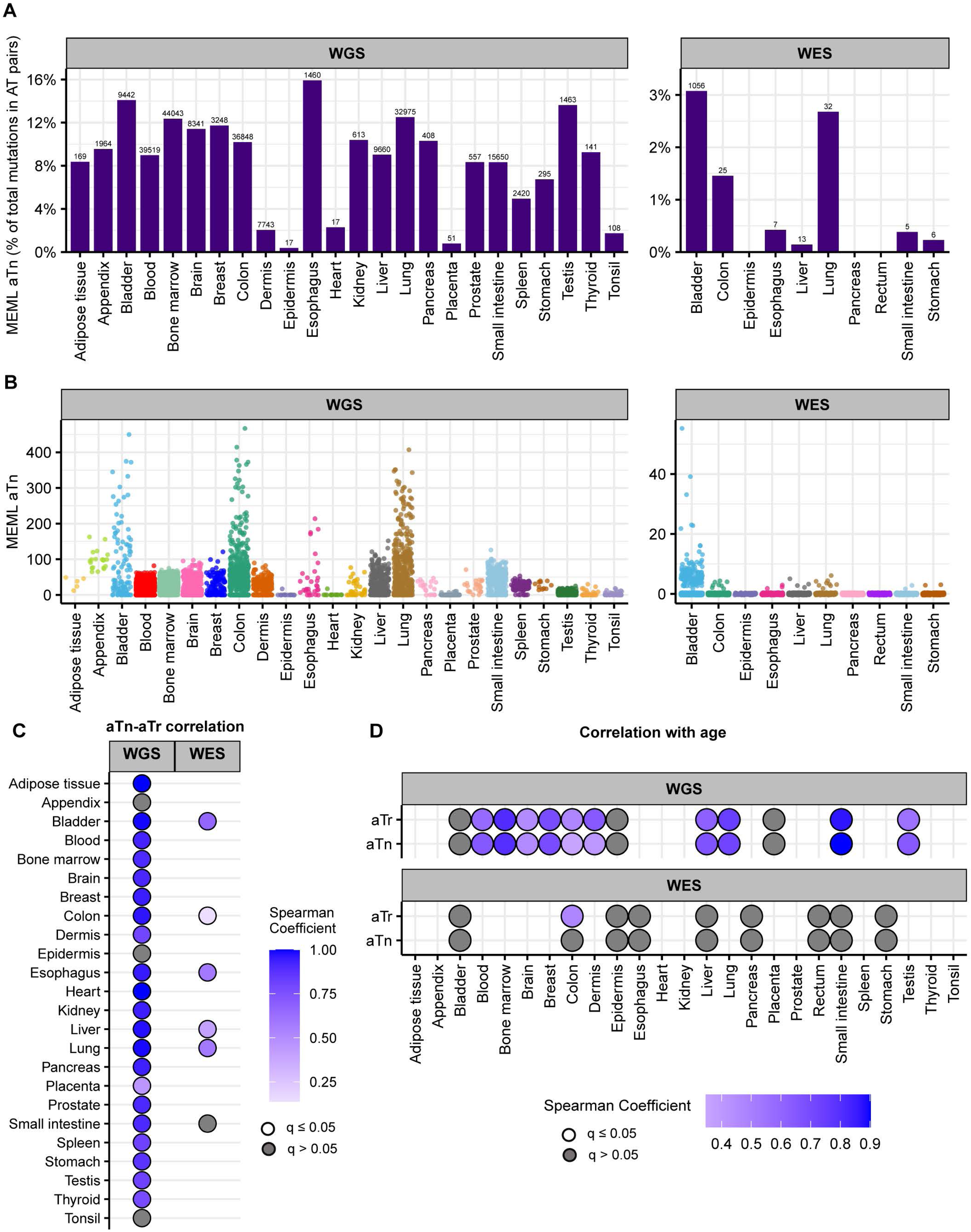
aTn mutational motif in healthy normal cells. **A**. The fraction (%) of aTn→aCn MEML within total AT-pair mutations (T→A, T→C, T→G, and reverse complement) is shown on Y-axis. The tissue types are plotted on X-axis. Left panel – WGS samples; right panel – WES samples. The total aTn MEML count for a tissue is shown above the respective bars. **B**. Sample-specific aTn→aCn MEML counts shown in jitter plots. X-axis indicates tissue types; labels above panels indicate WGS and WES groups. **C**. Correlation matrix showing aTn→aCn MEML versus aTr→aCr MEML to identify true aTn mutational motif. Spearman correlations were performed between sample-specific aTn and aTr MEML counts, including MEML=0 for either motif. Grey circles indicate P value >0.05 after correction for multiple hypothesis using Benjamini-Hochberg method within each sequencing group (WGS or WES). No circles indicate insufficient samples with MEML>0 for a tissue type to perform correlation analyses. **D.** Correlation matrix showing donor age versus mean donor MEML of aTn and aTr motifs. Spearman correlations were performed between donor age and mean MEML count of all samples from an individual donor for the indicated tissues and mutational motifs, including MEML=0 for both motifs. Grey circles indicate P value >0.05 after correction for multiple hypothesis using Benjamini-Hochberg method for a given motif within each sequencing group (WGS or WES). No circles indicate insufficient individual donors with mean MEML>0 for to perform correlation analyses. All source data and statistical analyses can be found in Supplementary Table S5.

T→C substitutions are central to aTn motif, however other mutagenic processes causing the same substitutions can confound statistical evaluation of aTn enrichment. Another motif centered around T→C, nTt→nCt (nTt in abbreviated format) is generated by the error-prone trans-lesion synthesis (TLS) across UV-induced cyclo-pyrimidine dimers (CPDs) formed by two adjacent thymines. It was revealed as a secondary minor UV-associated motif (40, 47, 55). Importantly, aTn motif is a part of nTt motif and vice a versa – i.e. the motifs overlap (See Supplementary Table S2). Such an overlap would not allow an unambiguous assignment of mutagenic mechanism, UV or S_N_2 reacting electrophiles, because of mutually confounding effects of respective diagnostic enrichment and MEML estimates of aTn and nTt motifs. This difficulty can be resolved using non-overlapping parts (i.e., sub-motifs) of mutually confounding motifs (Supplementary Table S2). To identify the presence of bona fide aTn contribution into MEML, we performed Spearman correlation tests between MEML counts of the aTn and its sub-motif, aTr (Supplementary Table S5c). In the aTr motif, thymine is followed by either an adenine or a guanine which eliminates any contribution of UV-mediated thymine-thymine dimers of the nTt motif. Significant positive correlation was detected between aTn and aTr MEML across all tissue types except appendix, epidermis, and tonsil (Fig. 4C, Supplementary Fig. S5, S6A). Lack of significant aTn-aTr correlation in epidermis and tonsil can be attributed to the low number of samples with both aTn and aTr MEML (Supplementary Fig. S5, S6A), however confounding effect of other mutagenic mechanisms cannot be excluded, especially in epidermis where UV mutagenesis could occur. Correlation analyses excluding either aTn MEML=0 or aTr MEML=0 further confirms that the strong contribution of aTn MEML in most tissues cannot be explained by confounding UV-mutagenesis (Supplementary Fig. S6A, Supplementary Table S5c).

Next, we asked if the aTn MEML accumulation shows clock-like feature across normal human tissues as previously identified in cancers and in healthy skin (41). We calculated mean MEML load in each donor and performed spearman correlation analyses with donor age within each tissue (Supplementary Table S5d). We detected significant positive correlation with age in all tissues where aTn and aTr was detected, except in bladder (Fig. 4D, Supplementary Table S5e). Of note, aTr shows a stronger correlation with age than aTn in dermis, suggesting a UV-like component (nTt) that is not age dependent. Correlation analyses between donor age and aTn or aTr MEML excluding zero MEML values reproduced same finding (Supplementary Fig. S6B). Together, our analyses showed similar features of aTn motif in healthy tissues as detected in tumors and generalized its’ clock-like feature to include multiple tissue types.

### Increased prevalence of aTn→aCn motif in various noncancer diseases

Next, we sought to compare the prevalence of the knowledge-based motifs in the healthy tissues with the diseased tissue cohorts, which included samples from 6 tissue types and 15 non-cancerous disease categories. We detected aTn motif MEML in brain, colon, liver, lung, and small intestine diseased samples (Fig. 2B, Supplementary Table S6a). Because aTn is an age-dependent motif, we categorized our cohort into three tertiles (0-33.3, 33.4-66.7, 66.8-100 years) of age groups to minimize the confounding effect of age while comparing mean aTn MEML of healthy and diseased donors using Wilcoxon Rank Sum test. We also only compared tissues originating from the same study within each age group to further minimize batch effect (Fig. 5, Supplementary Table S6b), and corrected p values for multiple hypothesis testing. To further account for the effect of age in different disease conditions, we performed multivariable regression analyses including age and diseases as predictor variables (Supplementary Fig. S7, Supplementary Table S6c). We also performed Spearman correlation analysis between donor age and mean donor MEML to detect clock-like accumulation of aTn MEML in diseased tissues (Supplementary Table S6d).

**Figure 5:**
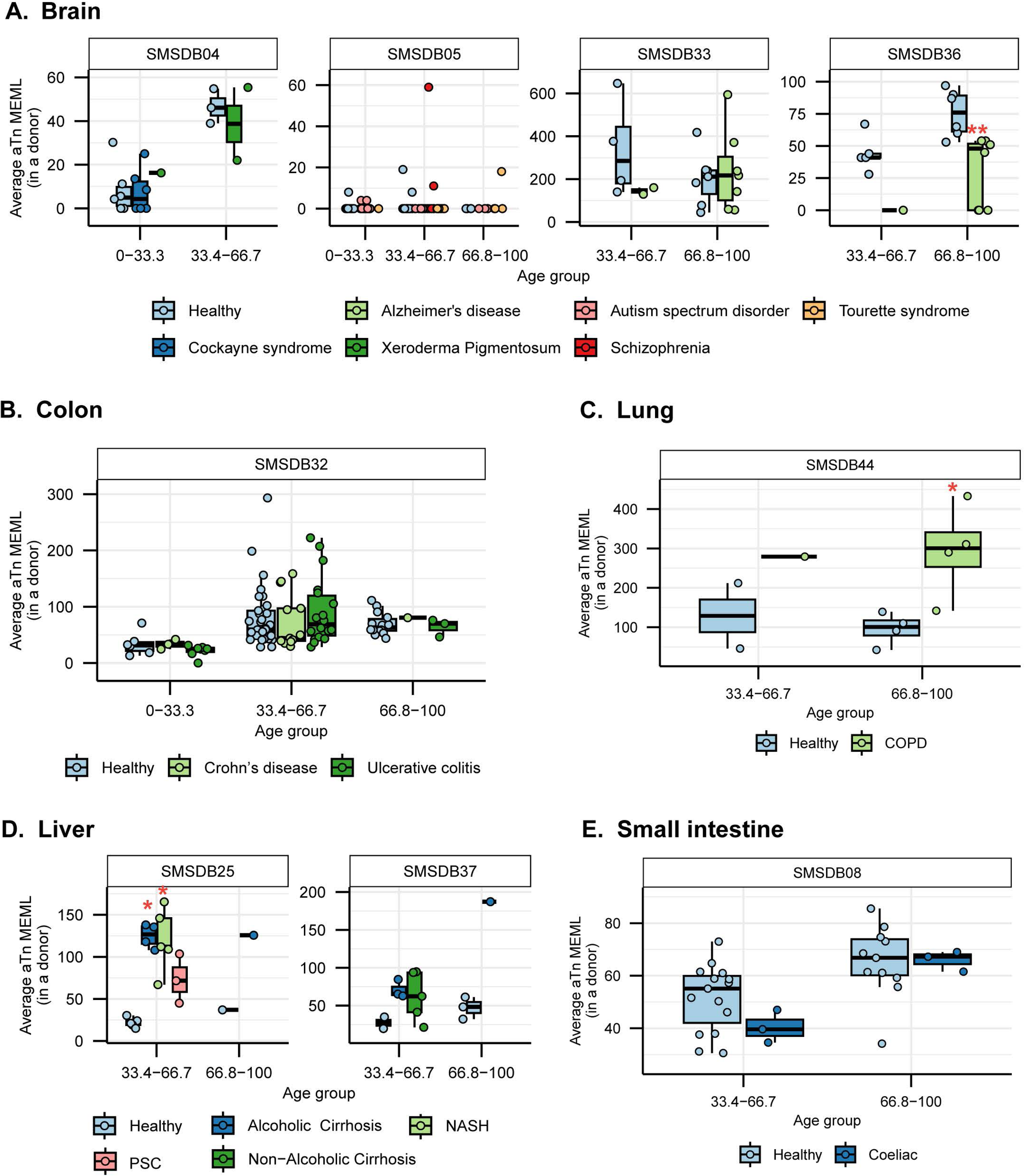
aTn motif in diseased tissues. Mean donor aTn MEML for healthy and diseased samples are shown in a boxplot for brain **(A**), colon **(B)**, lung **(C)**, liver **(D)**, and small intestine **(E)**. X-axes indicate age groups where samples and donors were available. Labels above panels indicate source study of the samples. Analyses of only WGS samples are shown. Asterisks in boxplots represent statistical significance (Wilcoxon Rank Sum test) compared to the healthy donors within each study and age group after correcting for multiple hypothesis testing using Benjamini-Hochberg method. *P value ≤ 0.05, **P value ≤ 0.01, ***P value ≤ 0.001. All source data and statistical analyses can be found in Supplementary Table S6.

We did not see any statistically significant difference between healthy and diseased cohorts in the first tercile (0-33.33) of age group, regardless of the disease type. In brain, we detected statistically significant reduction of aTn MEML in donors with Alzheimer’s disease (Fig. 5A). Multivariable regression analyses without age categorization identified statistically significant positive effect of Alzheimer’s disease on aTn MEML load, while Tourette syndrome showed significant negative effect (Supplementary Fig. S7A). Spearman correlation analyses did not reveal age dependency of aTn accumulation in any brain diseases (Supplementary Table S6d).

In colon, we did not identify any significant association of aTn MEML with disease conditions in either age-matched Mann-Whitney test (Fig. 5B) or multivariable regression analyses without age grouping (Supplementary Fig. S7B). Spearman correlation analyses detected age dependent accumulation of aTn MEML in both Crohn’s (Spearman’s *ρ*= 0.5, q=0.03) and ulcerative colitis (Spearman’s *ρ*= 0.6, q=<0.001) samples (Supplementary Table S6d), suggesting steady, continuous accumulation of aTn motif mutations in these diseased tissues over donor lifetime.

Lung sample donors with Chronic Obstructive Pulmonary Disorder (COPD) showed a clear disease-dependent accumulation of aTn MEML (Fig. 5C). Compared to healthy donors, donors with COPD showed a 3-fold increase in the average aTn MEML. Multivariable regression analyses identified strong positive association of COPD with aTn MEML after accounting for the effect of age (Supplementary Fig. S7C), while no clock-like activity was identified by Spearman correlation analyses (Supplementary Table S6d).

High aTn MEML was detected across liver tissue donors with alcoholic cirrhosis and Non-Alcoholic Steatohepatitis (NASH) compared to healthy donors (Fig. 5D). Donors with Primary Sclerosing Cholangitis (PSC) also showed increased aTn mutagenesis but were not significantly associated after correcting for multiple hypothesis (Supplementary Table S6b). Multivariable regression analysis without age categorization showed strong positive association of all liver diseases except non-alcoholic cirrhosis, while the effect of age was not significant (Supplementary Fig. S7D). Only donors with NASH (n=5) showed a strong age-dependent accumulation of aTn MEML (Spearman’s *ρ*= 0.9, q=0.004) (Supplementary Table S6d).

Small intestine sample donors with coeliac disease did not show any association with aTn MEML (Fig. 5E). Regression analysis identified age as the strongest predictor of aTn MEML accumulation in small intestines, with coeliac disease showing a negative but statistically insignificant association (Supplementary Fig. S7E). No significant clock-like aTn accumulation was detected by correlation analyses in samples with coeliac disease (Supplementary Table S6d).

Together, these findings suggest that diseases associated with heightened exposure to exogenous DNA damaging agents such as tobacco smoke in COPD and alcohol in cirrhotic liver disease exhibit significantly elevated aTn mutagenesis, consistent with increased S_N_2 electrophile burden driven by xenobiotic metabolism and its downstream mutagenic consequences.

### Clock-like mutagenesis caused by deamination of 5-methylcytosine motif nCg→nTg is highly enriched in multiple human tissues

In mammalian genomes, 70%-80% of the cytosines in CpG dinucleotides (nCg motifs) are methylated, and spontaneous 5-methylcytosine deamination results in C→T substitutions (56). This nCg motif mutagenesis is ubiquitously detected across all tumor types in WES and WGS TCGA the PCAWG catalogs, shows clock-like accumulation, and corresponds with COSMIC signature SBS1 (3, 18).

In our cohort of WGS samples in healthy tissues, we detected nCg MEML across all tissue types with 14.15% median nCg→nTg MEML contribution in total CG-pair mutations (Fig. 6A). WES samples showed 15.5% median nCg contribution in CG-pair mutations (Fig. 6A) (Supplementary Table S6a). Samples with nCg MEML were detected across all tissue types with sample-specific MEML ranging from zero to 2,010 in WGS samples, and from zero to 1,456 in WES samples (Fig. 6B, Supplementary Table S7b).

**Figure 6:**
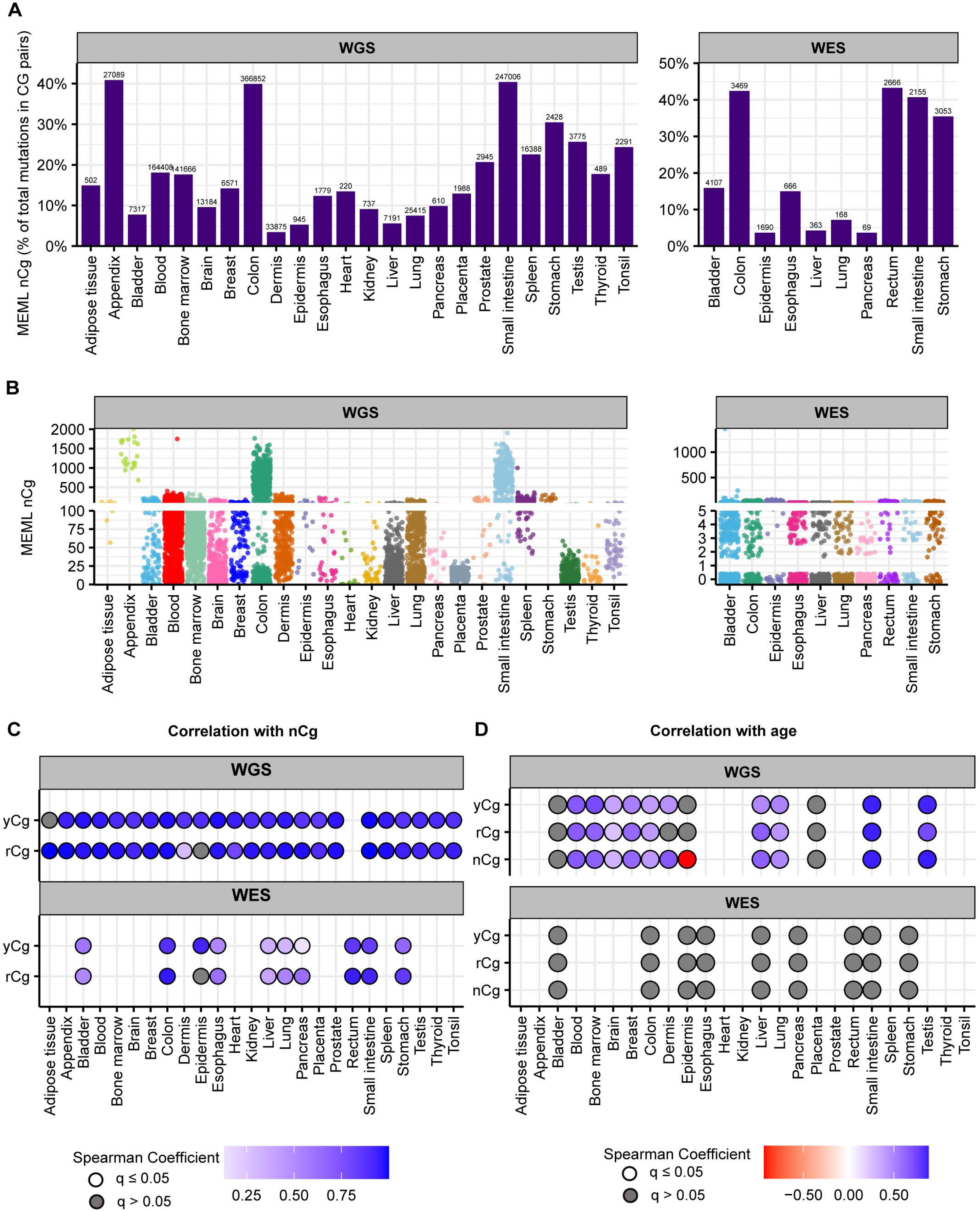
nCg mutational motif in healthy normal cells. **A.** The fraction (%) of nCg→nTg MEML count within total CG-pair mutations (C→A, C→G, C→T, and reverse complement) is shown on Y-axis. The tissue types are plotted on X-axis. Left panel – WGS samples; right panel – WES samples. The total nCg MEML count for a tissue is shown above the respective bars. **B.** Sample-specific nCg→nTg MEML counts shown in jitter plots. X-axis indicates tissue types; labels above panels indicate WGS and WES groups. **C.** Correlation matrix showing nCg→nTg MEML versus rCg→rTg MEML, and nCg→nTg MEML versus yCg→yTg MEML, to identify true nCg mutational motif. Spearman correlations were performed between sample-specific nCg, rCg, and yCg MEML counts, including MEML=0 for either motif. Grey circles indicate P value >0.05 after correction for multiple hypothesis using Benjamini-Hochberg method within each sequencing group (WGS or WES). No circles indicate insufficient samples with MEML>0 for a tissue type to perform correlation analyses. **D.** Correlation matrix showing donor age versus mean donor MEML of nCg, rCg, and yCg motifs. Spearman correlations were performed between donor age and mean MEML count of all samples from an individual donor for the indicated tissues and mutational motifs, including MEML=0 for both motifs. Grey circles indicate P value >0.05 after correction for multiple hypothesis using Benjamini-Hochberg method for a given motif within each sequencing group (WGS or WES). No circles indicate insufficient individual donors with mean MEML>0 for to perform correlation analyses. All source data and statistical analyses can be found in Supplementary Table S7.

Multiple mutagenic processes cause cytosine deamination in mammalian tissues including CpG deamination, repair of UV-mediated cyclobutane pyrimidine dimers (CPDs), and activities of cytidine deaminases like AID and APOBEC (2). The trinucleotide base composition in nCg motif overlaps with base preference by UV mutagenesis, which forms pyrimidine-pyrimidine dimers in yCn context. To separate the contribution of yCn and detect true nCg MEML in tissues, we used the approach of sub-motif analyses and split the nCg motif into two sub-motifs, rCg and yCg (Supplementary Table S2). The mutable cytosine in rCg is preceded and followed by a purine, therefore excluding the possibility of CPD formation by UV-damage. yCg on the other hand has a pyrimidine preceding the mutable cytosine and can be converted to a thymine by both spontaneous and UV-mediated deamination. Performing Spearman correlation analyses of nCg MEML with both rCg MEML and yCg MEML allowed us to identify tissues that do not show true spontaneous CpG deamination (Supplementary Table S7c). We detected strong positive nCg-rCg and nCg-yCg correlations in all tissues and sequencing groups, except in dermis and epidermis where nCg-rCg correlation is absent (in epidermis) or weaker (in dermis) compared to nCg-yCg correlation (Fig. 6C, Supplementary Fig. S8A, S9A).

Correlation analyses excluding samples that had either motif MEML=0 showed strong nCg-rCg and nCg-yCg correlations in both epidermis and dermis samples (Supplementary Fig. S8B). To further dissect the effect of UV in skin samples, we analyzed the yCh→yTh motif which overlaps with UV-preferred yCn motif but excludes CpG motifs (Supplementary Tables S7b, S7c). We found strong nCg-yCh correlation in both dermis and epidermis samples (Supplementary Table S6c). Based on these sub-motif analyses we concluded that the spontaneous deamination of meCpG is detectable but can be heavily confounded by UV-mutagenesis in skin samples.

Next, we investigated the clock-like feature of meCpG deamination as previously detected in cancers (3). For donors with known age, we calculated average donor-specific MEML value for nCg motif and its sub-motifs, yCg and rCg (Supplementary Table S7d). Spearman correlation analyses showed significant positive correlation between donor age and all three motifs (nCg→nTg, rCg→rTg, yCg→yTg) in tissues where true spontaneous meCpG deamination activity was indicated by sub-motif analyses described above (Fig. 6D, Supplementary Fig. S9B, Supplementary Table S7e).

### Increased prevalence of nCg→nTg motif in various noncancer diseases

We detected nCg MEML in all disease categories across brain, colon, epidermis, liver, lung, and small intestine tissues (Fig. 2B, Supplementary Table S8a). As previously described for healthy samples, we performed spearman correlation analyses between nCg, rCg, and yCg MEML to identify true nCg contribution (Supplementary Table S8b). We detected strong positive correlation of nCg MEML with both rCg and yCg MEML in all disease categories except psoriasis, a skin disease, which only showed correlation with yCg (Supplementary Fig. S10). We excluded psoriasis from further analyses of association with nCg motif.

Association of diseases with nCg motif MEML showed a similar pattern as that with aTn motif MEML. Donors with Alzheimer’s disease showed a reduction in nCg motif accumulation (Fig. 7A), while donors with COPD showed significantly increased nCg MEML accumulation (Fig. 7C). Donors with alcoholic cirrhosis, NASH, and PSC showed higher nCg MEML with PSC association being statistically insignificant after FDR correction (Fig. 7D, Supplementary Table S8c). Small intestine sample donors with coeliac disease also did not show any significant difference in nCg MEML compared to healthy donors (Fig. 7E). Contrary to aTn MEML, statistically significant increase of nCg MEML was detected in donors with ulcerative colitis in the first age terciles (Fig. 7B).

**Figure 7:**
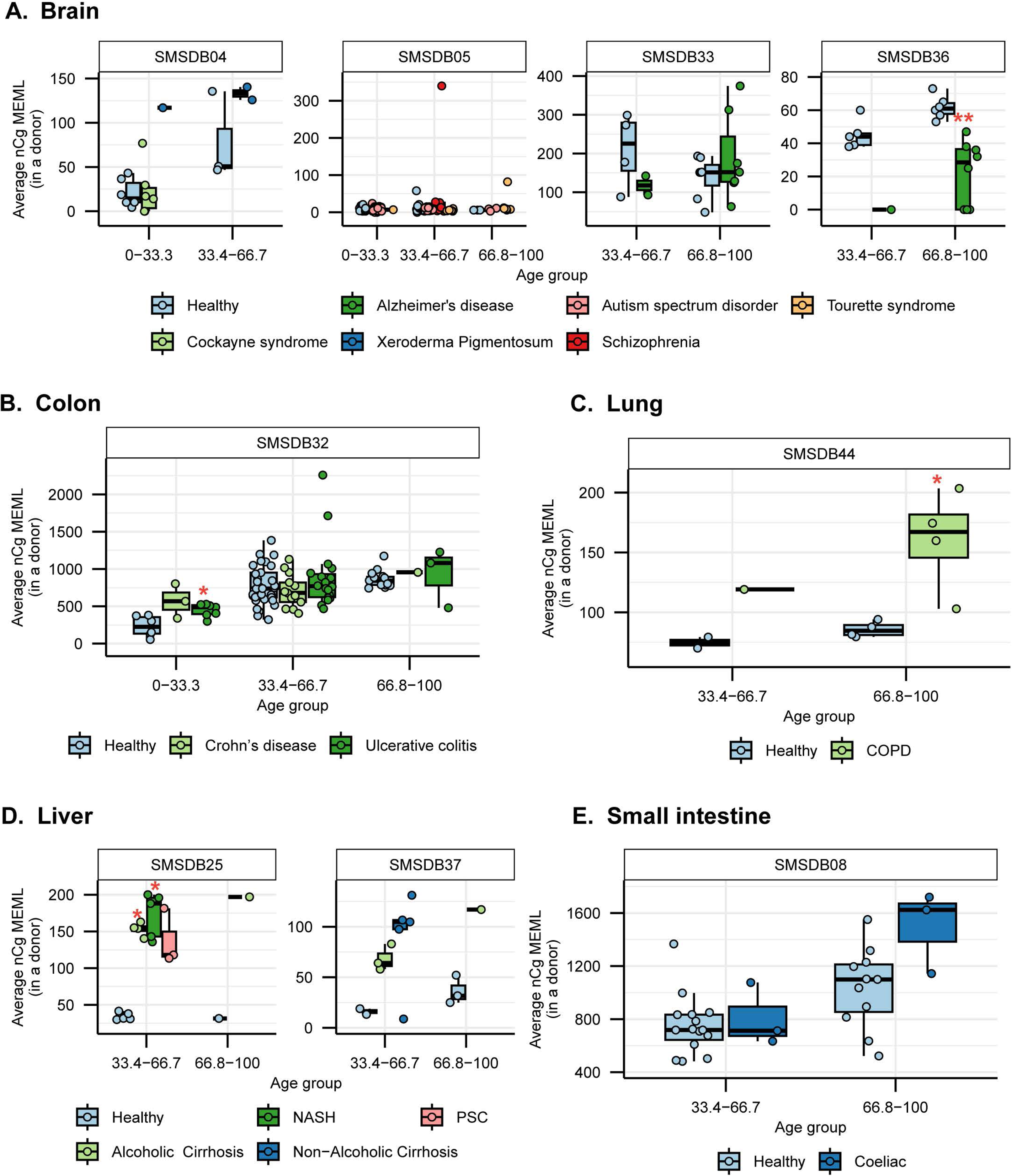
nCg motif in diseased tissues. Mean donor nCg MEML for healthy and diseased samples are shown in a boxplot for brain **(A**), colon **(B)**, lung **(C)**, liver **(D)**, and small intestine **(E)**. X-axes indicate age groups where samples and donors were available. Labels above panels indicate source study of the samples. Analyses of only WGS samples are shown. Asterisks in boxplots represent statistical significance (Wilcoxon Rank Sum test) compared to the healthy donors within each study and age group after correcting for multiple hypothesis testing using Benjamini-Hochberg method. *P value ≤ 0.05, **P value ≤ 0.01, ***P value ≤ 0.001. All source data and statistical analyses can be found in Supplementary Table S8.

Multivariable regression analyses after adjusting for the effect of age showed significant positive association of increased nCg MEML with Xeroderma Pigmentosum and Alzheimer’s disease in brain, ulcerative colitis in colon, COPD in lung, NASH, PSC, alcoholic and non-alcoholic cirrhosis in liver, and coeliac disease in small intestine tissues (Supplementary Fig. S7, Supplementary Table S8d). Spearman correlation detected clock-like accumulation of nCg MEML only in donors with Tourette’s Syndrome and ulcerative colitis (Supplementary Table S8e).

Together, the similar trend of both aTn and nCg MEML accumulation in inflammatory diseases suggests dysregulation of endogenous mutagenic processes by disease-specific physiologies.

### Motifs associated with exogenous and metabolic chemical exposures reveal sporadic tissue- and disease-specific mutagenesis

We detected sporadic enrichment of five knowledge-based motifs aCy, cTg, gCn, hTg, and tgC (see mechanistic details in Table 1) associated with exposures to exogenous and metabolic stress-induced chemicals across healthy and diseased tissue types (Supplementary Fig. S11, Supplementary Table S9a). In healthy tissues, enrichment of these motifs was detected in a small fraction of samples across multiple tissue types, with gCn motif associated with acetaldehyde exposure showing the broadest tissue distribution (Supplementary Fig. S11A). hTg motif associated with exposure to MMS, an S_N_2-reacting chemical, was detected in only a fraction of samples and tissues compared to aTn motif, further supporting aTn-specificity by a broader suite of S_N_2-reacting chemicals. We did not detect clock-like property for any of these motifs, as is expected for exposure-dependent mutagenesis (Supplementary Table S9b).

In diseased tissues, we detected tissue-specific elevation of some motifs in association with specific disease conditions (Supplementary Table S9c). tgC motif associated with redox stress were not detectable in small intestine samples of healthy donors but were detectable in few donors with coeliac disease (Supplementary Fig. S11B). hTg motif, similar to aTn motif, showed increased mutagenesis in lungs of donors with COPD (Supplementary Fig. S11C). Donors with COPD also showed elevated mutagenesis of acetaldehyde-associated gCn motif and cTg motif associated with exposure to aristolochic acid (Supplementary Fig. S11D, E). gCn motif MEML was also elevated in colon samples of donors with ulcerative colitis (Supplementary Fig. S11D).

Together, the sporadic prevalence of these chemical exposure motifs in healthy tissues and their tissue-specific elevation in select disease conditions suggest that mutagenesis by exogenous and endogenous reactive chemicals, while less ubiquitous than the clock-like aTn and nCg motif-associated processes, contributes to the somatic mutation burden of specific tissues and may be exacerbated by disease-associated physiological perturbations.

### Mutation motifs yCn→yTn and nTt→nCt detected in several tissues can support UV mutagenesis only in skin

UV radiation is a common and potent environmental mutagen that facilitates many types of genome changes including SBSes. UV causes covalent bond formation between two adjacent pyrimidines that results in cyclobutane pyrimidine dimers (CPD) and pyrimidine 6–4 pyrimidone (6-4PP) formation, with CPDs contributing to >80% of the UVB-mediated mutations in mammalian cells (55). C→T mutations in the yCn context are the most frequent result of UV lesions. The yCn→yTn mutations can result from direct error-prone copying of a cytosine within a CPD by TLS polymerases. Importantly, spontaneous deamination of cytosine changing it to uracil is increased several orders of magnitude in CPDs (65) and can result in C→T substitutions after accurate copying by TLS polymerases and another round of replication. Less frequently, thymine in T-T CPDs can be copied by TLS polymerases causing T→C mutations in the nTt context. While less frequent, nTt motif mutations correlate with yCn in normal human skin and in cancers (40, 47).

We detected yCn MEML in dermis and epidermis, where UV-mutagenesis is expected, as well as in bladder, blood, lung, and spleen (Fig. 8A, B, Supplementary Table S10a, 10b). Dermis samples show a high contribution of yCn MEML in CG-pair mutations, second only to epidermis samples (Fig 8A, Supplementary Table S10a). To independently verify UV mutagenesis in tissues where yCn MEML was detected, we performed spearman correlation analyses of yCn MEML with nTt MEML (a minor motif for UV-mutagenesis) and with rCg MEML (a confounding sub-motif of nCg mutagenesis in meCpG) (Supplementary Tables S2 and S10c). There was significant positive yCn-nTt correlation in skin samples and blood, while yCn-rCg significant positive correlation was detected in lung and bladder (Fig. 8C). Removal of samples that does not have a MEML count for either motif showed a significant positive yCn-nTt correlation only in dermis (WGS) and epidermis (WES) (Fig. 8D). Based on the sub-motif analyses we concluded that only skin samples show UV-mediated yCn mutational motif; the source of yCn MEML in other tissues can likely be attributed to spontaneous CpG deamination or other mutagenic processes. We further calculated donor-specific yCn and nTt MEML values for dermis and epidermis (Supplementary Table S10d) and assessed correlation with donor age (Supplementary Table S10e). We do not see age-dependent accumulation of UV mutational motifs in skin, except for negative correlation between nTt MEML and donor age in epidermis (Fig. 8E, Supplementary Table S10e). The total mutation burden in these donors also showed a negative trend of detection with donor age (Spearman’s *ρ* −0.67, p-value 0.06). Since there was only small number of donors and epidermis samples with non-zero nTt MEML, this correlation should be reconfirmed by independent studies.

**Figure 8:**
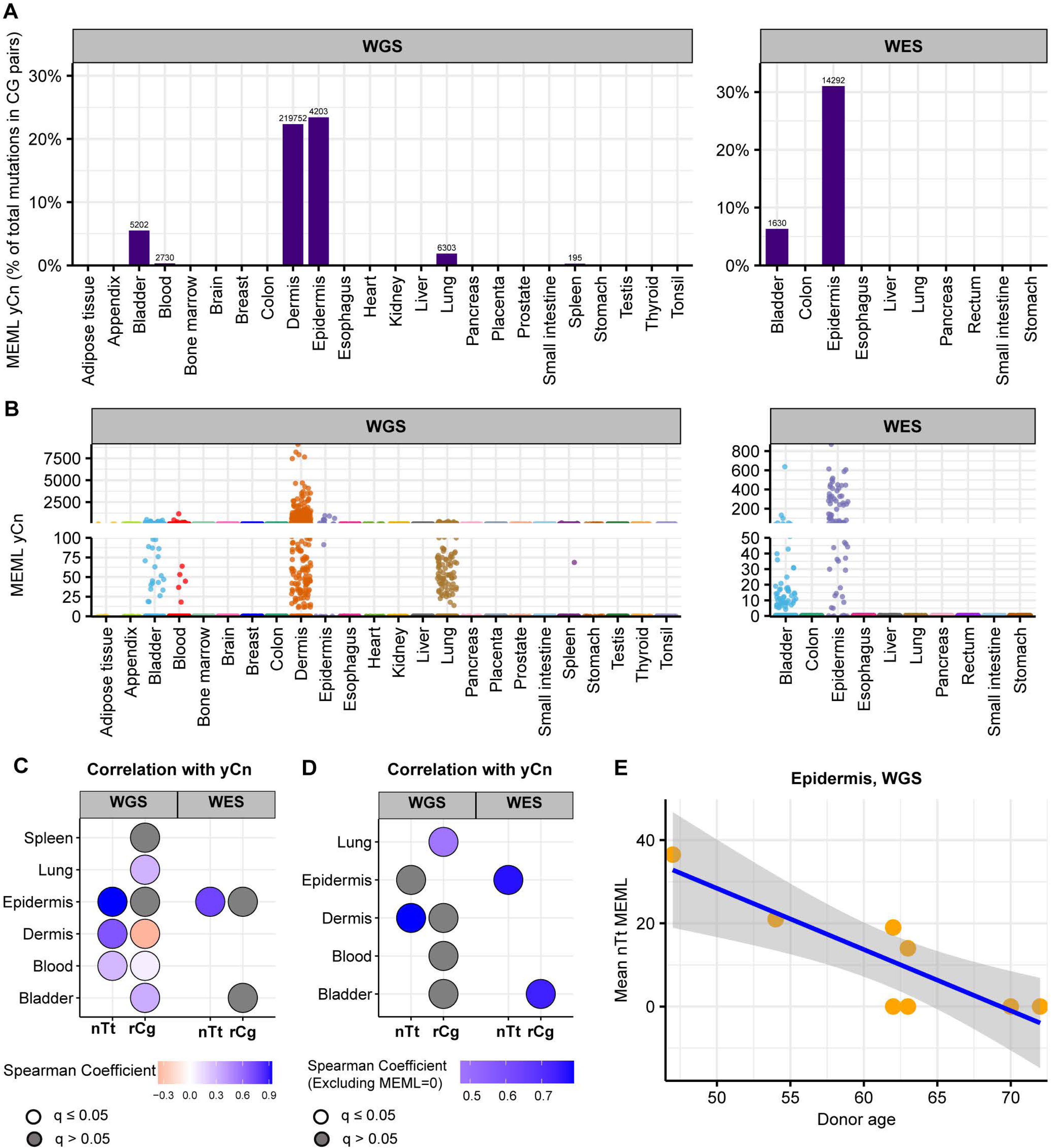
UV mutational motifs in healthy normal cells. **A.** The fraction (%) of yCn→yTn MEML count in total CG-pair mutations (C→A, C→G, C→T, and reverse complement) is shown on Y-axis. The tissue types are plotted on X-axis. Left panel – WGS samples; right panel – WES samples. The total yCn MEML count for a tissue is shown above the respective bars. **B.** Sample-specific yCn→yTn MEML counts shown in jitter plots. X-axis indicates tissue types; labels above panels indicate WGS and WES groups. **C.** Spearman correlation matrix showing correlations of sample-specific values of yCn→yTn MEML versus nTt→nCt MEML, and yCn→yTn MEML versus rCg→rTg MEML to eliminate confounding effects of overlapping motifs and to support hypothesis about minor UV-motif contribution of UV-mutagenesis in tissue groups that contained samples with yCn MEML. Grey circles indicate P value >0.05 after correction for multiple hypothesis using Benjamini-Hochberg method within each sequencing group (WGS or WES). No circles indicate insufficient samples with MEML>0 for a tissue type to perform correlation analyses. Samples with MEML=0 for either motif was included. **D**. Same as **C**, but samples with MEML=0 for either motif were excluded. **E.** Mean of nTt MEML values (including MEML=0) for samples from epidermis of each donor plotted against donor age. Only epidermis, but not dermis, showed statistically significant correlation after correction for multiple hypotheses testing (Spearman’s ρ= −0.8, q=0.03). All source data and statistical analyses for all tissue types can be found in Supplementary Table S10.

### Motif analyses confirm mutagenesis by APOBEC3A and APOBEC3B in bladder, breast, lung, small intestine, and liver

APOBEC (apolipoprotein B mRNA-editing enzyme, catalytic polypeptide-like) cytidine deaminases, part of the antiviral innate immunity and RNA editing systems, are also known to cause hypermutation in genomes of human cancers (2, 52, 64, 66). APOBECs cause Cytosine (C) to Uracil (U) deamination in single-stranded DNA, which can then lead to different base substitution outcomes contingent on processing by the uracil-DNA glycosylases. If no glycosylation occurs, the replicative polymerases can accurately copy the deaminated uracils as thymines and generate C→T mutations. If uracils are cleaved by glycosylation, the resulting AP-sites sites are copied by error-prone TLS polymerases which can lead to either C→T, C→G, or C→A mutations ((67, 68) and references therein). Analyses of large cancer datasets have shown over representation of APOBEC-mediated C→T and C→G mutations in tCw context, with C→A mutations being barely distinguishable from random mutagenesis (52, 64). The tCw→tTw mutational motif closely resembles COSMIC signature SBS2, while tCw→tGw is analogous to SBS13 (3, 18, 52). APOBEC-induced tCw mutational motif enrichment, as well as mutation clusters (also referred as kataegis) have been identified in multiple cancer types (18, 52, 63, 66, 69). Signature analyses have also identified APOBEC mutagenesis in several healthy tissues (16, 17, 49).

We detected enrichment of tCw mutational motif across several tissues in our dataset (Fig 9A-9C, Supplementary tables 11a, 11b). tCw MEML (including tCw→tTw and tCw→tGw) contributed up to 11% of total CG-pair mutations in the WGS samples and up to 7% in WES samples (Fig. 9A, Supplementary Table S11a). APOBEC-mediated mutations tend to be localized in regions of ssDNA stretches, resulting in clusters of strand-coordinated G or C mutations with inter-mutation distance of 10-10^4^ bases (63, 64, 66, 70). To further granulate the features of APOBEC mutagenesis, we identified the load and pattern of clustered mutations in our dataset. The percentage of samples with >0 mutation clusters ranged 12%-100% in WGS samples and 4%-19% in WES samples (Supplementary Table S11c). We then isolated the samples with total cluster count >0 and calculated the mean number of clusters, stratified by tissue and cluster type. We detected a high bias towards non-coordinated clusters with contributions of both G- or C-coordinated and of A- or T-coordinated clusters across many tissues (Fig. 9D, Supplementary Table S11d).

**Figure 9:**
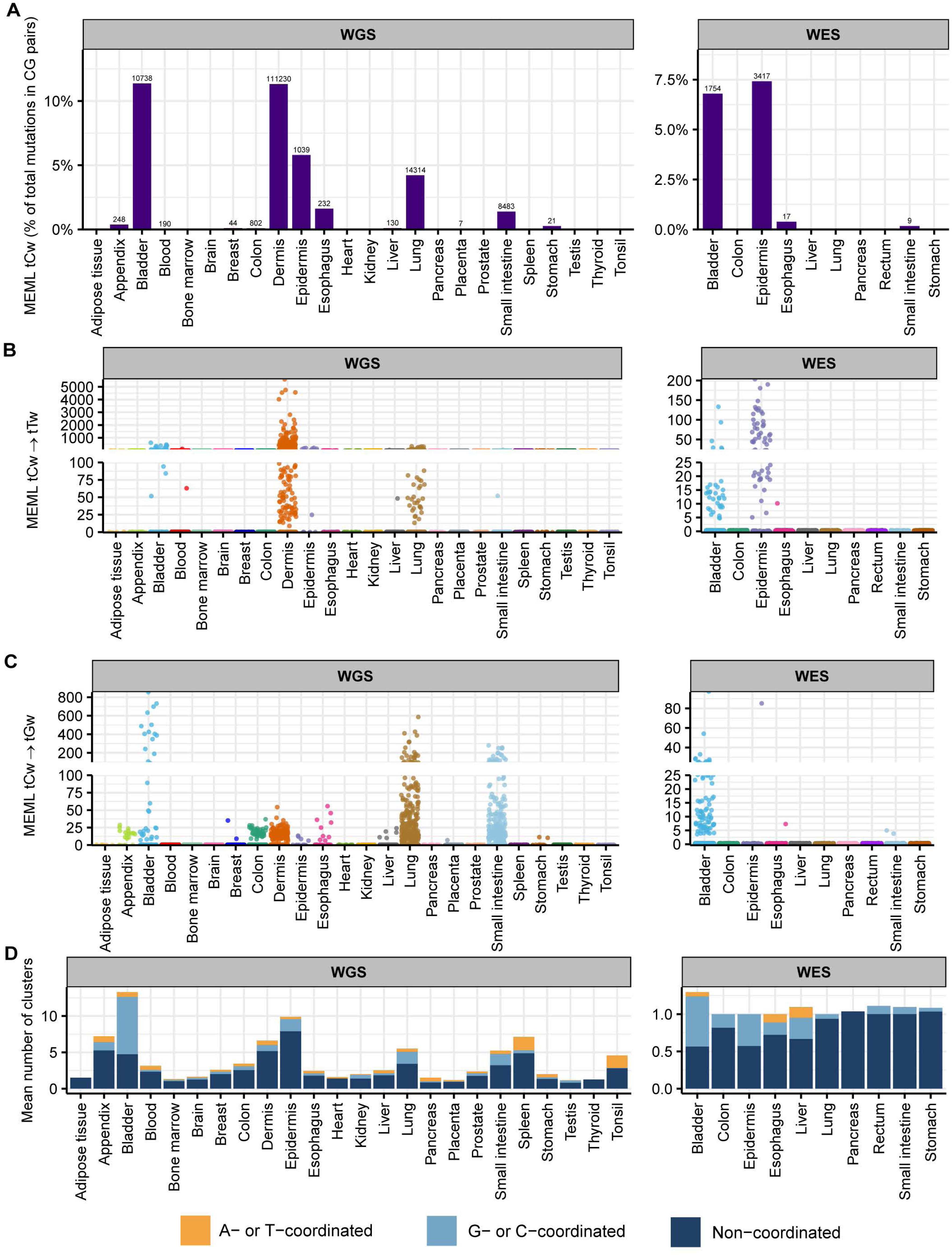
APOBEC tCw mutational motif in healthy normal cells. **A.** The fraction (%) of tCw→tTw and tCw→tGw MEML count in total CG-pair mutations (C→A, C→G, C→T, and reverse complement) is shown on Y-axis. The tissue types are plotted on X-axis. Left panel – WGS samples; right panel – WES samples. The total tCw MEML count for a tissue is shown above the respective bars. **B.** Sample-specific tCw→tTw MEML counts shown in jitter plots. X-axis indicates tissue types; labels above panels indicate WGS and WES groups. **C.** Sample-specific tCw→tGw MEML counts shown in jitter plots. X-axis indicates tissue types; labels above panels indicate WGS and WES groups. **D.** Mean number of mutation clusters for samples with total number of clusters >0 in a tissue type. All source data and statistical analyses can be found in Supplementary Table S11.

To refine tissues with detectable contribution of APOBEC mutagenesis into mutation load from tissues where tCw MEML was detected, we defined three specific features as indicators of APOBEC mutagenesis: (i) presence of samples with statistically significant enrichment of combined tCw→tTw and tCw→tGw APOBEC motifs (Supplementary Fig. S12A), (ii) high prevalence of both tCw→tTw and tCw→tGw within G- or C-coordinated clusters in samples showing enrichment with combined APOBEC motif (Supplementary Fig. S12B), and (iii) high prevalence of G- or C-coordinated clusters (Supplementary Fig. S12C). In Supplementary Figure S8, we present additional plots for tissues that satisfied criteria for APOBEC mutagenesis. We identified multiple samples in bladder, breast, lung, liver, and small intestine tissues that showed APOBEC enrichment (Supplementary Fig. S12A). In agreement with prior knowledge about APOBEC mutagenesis, there were comparable numbers of tCw→tTw and tCw→tGw mutations in G- or C-coordinated clusters of these tissues, with a small contribution of tCw→tAw mutations (Supplementary Fig. S12B). APOBEC enrichment is expected to be less in smaller clusters and scattered mutations due to a higher chance of these clusters being formed by random mutagenesis in close vicinity. We detected APOBEC enrichment in G- or C-coordinated clusters of different sizes across all five tissues (Supplementary Fig. S12C). While some of these features were also detectable in dermis samples, APOBEC mutagenesis in G- or C-coordinated clusters was heavily skewed towards tCw→tTw MEML (Supplementary Data S1). Since tCw motif overlaps with UV-preferred yCn motif and causes the same base substitution, we could not distinguish between the contribution of APOBEC and UV mutagenesis in skin samples. In summary, motif-centered analyses lead us to conclude that APOBEC mutagenesis is detectable in samples of normal bladder, breast, liver, lung, and small intestine tissues.

Next, we sought to identify the preference for APOBEC3A (A3A) or APOBEC3B (A3B) in tissues where APOBEC mutagenesis was identified. The A3A have been identified as greater source of cancer mutators, however contribution of A3B is also detectable (36, 38, 52). Experiments in yeast model system, and subsequent analyses of cancer datasets have revealed that hypermutation motif preference is distinguishable between A3A and A3B (36). While A3A prefers ytCa motif (APOBEC-preferred tCa motif preceded by a pyrimidine), A3B prefers rtCa motif (APOBEC-preferred tCa motif preceded by a purine). We calculated the enrichment of both ytCa (A3A-like) and rtCa (A3B-like) motifs across different mutation categories in bladder, breast, liver, lung, and small intestine tissues (Supplementary Fig. S12D, Supplementary Table S11e). Where enrichment of both ytCa and rtCa were statistically significant (Fisher’s exact test), we performed Breslow-Day test of homogeneity to compare the odds ratios of the two motif enrichments (Supplementary Table S11f). Bladder, breast, lung, and small intestine showed statistically significant higher ytCa (A3A-like) enrichment than rtCa (A3B-like) across all clustered mutations (Supplementary Fig. S12D). Bladder and lung also showed preference for ytCa (A3A-like) enrichment in scattered and genome-wide mutations. Liver showed a preference for rtCa (A3B-like) enrichment; however, it was not detected to be significantly higher than ytCa (A3A-like) enrichment as both enrichment levels were relatively low compared to the other tissues. Together, our analyses revealed A3A to be the predominant APOBEC mutator in bladder, lung, breast, and small intestine healthy tissue samples.

## Discussion

In this study, we performed a meta-analysis of the somatic mutation catalogues of cancer-free normal tissues generated in dozens of independent studies to find common sources of mutations. We formed stringent statistical hypotheses to identify the enrichment of expert-curated and experimentally validated mutational motifs preferentially mutated by known mutagenic processes in the mutation catalogues of individual samples. We identified the activities of some known mutagenic mechanisms in expected tissue types, like UV-mutagenesis in skin tissues and clock-like deamination of meCpG motifs across all tissue types. We also found new, ubiquitous motif aTn→aCn associated with exposure to small epoxides and other S_N_2 electrophiles across all tissues analyzed. Our previous study had identified this motif to be clock-life in multiple cancers across the PCAWG cancer catalogue (41). Here we show that similar to the spontaneous deamination of methylated CpGs, mutagenesis by exposure to small epoxides and potentially a broader class of S_N_2 electrophiles accumulates throughout human lifetime and predates diseased states. We also distinguished between the preference for APOBEC3A-like and APOBEC3B-like activity in five tissue types, bladder, breast, lung, liver, and small intestine, where mutagenesis by APOBEC deaminases was detected in this current study and previously reported for both normal and cancerous tissues of some of these organs. These findings were enabled by the application of mutational motif-centered pipeline P-MACD accounting for prior knowledge accumulated in mechanistic research revealing the motif preference of mutagens and the molecular mechanisms of mutagenesis. The following discussion outlines the composition and limitations of the general framework of bioinformatics and statistical analyses that can be utilized for elucidating contributions of endogenous and environmental mutagenic factors into base substitution loads in human tissues.

### Framework utilizing knowledge-based motif-centered analyses to dissect the sources of mutagenesis in somatic tissues of individuals

Knowledge produced by prior mechanistic research about distinct trinucleotide motif preference of a mutagenic factor to generate mutations was used for calculating statistically supported enrichment and MEML attributable to the motif-associated mutagenesis. There are only a few trinucleotide motifs so far that have been robustly associated with mutagenic factors (Table 1), but more motifs may be discovered from future efforts of the research community. Since there are only 4 bases and 96 trinucleotide motifs in DNA, a partial overlap, or even complete similarity between motifs preferred by different mutagens is inevitable. However, information derived from mechanistic research can also be used to identify non-overlapping components of motifs (i.e., sub-motifs) that are expected to be preferentially mutagenized by one out of two or more mechanisms associated with the overlapping motifs (step A, Fig. 10). Motifs and sub-motifs derived from mechanistic knowledge were used to perform motif-centered analyses and generate statistically supported enrichment and MEML values from the mutation catalogs of each individual sample (step B, Fig. 10). In downstream analyses (step C, Fig. 10), these partially interchangeable values were used to explore correlations between different motifs and/or sub-motifs. Revealed correlations and prior mechanistic knowledge were used to resolve ambiguity stemming from motif-motif overlaps. For example, we used correlation between aTn→aCn motif and its sub-motif aTr→aCr, which does not overlap with a minor UV-motif nTt→nCt, to dissect and rule out enrichment of epoxide mutagenesis in epidermis (Fig. 4C). Importantly, mutation load in epidermis was enriched for major UV-motif yCn→yTn, which in turn correlated with the minor UV-motif nTt→nCt (Fig. 8C, D). In general, step of motif and sub-motif construction is aided by correlation analysis applied to motif-motif pairs chosen based on sequence overlap and prior knowledge about mutagens underlying motifs. Sample-specific values produced by motif-centered pipeline can also be utilized in downstream statistical analyses to compare cohorts and perform correlations with the biological features of the samples and donors within cohorts.

**Figure 10:**
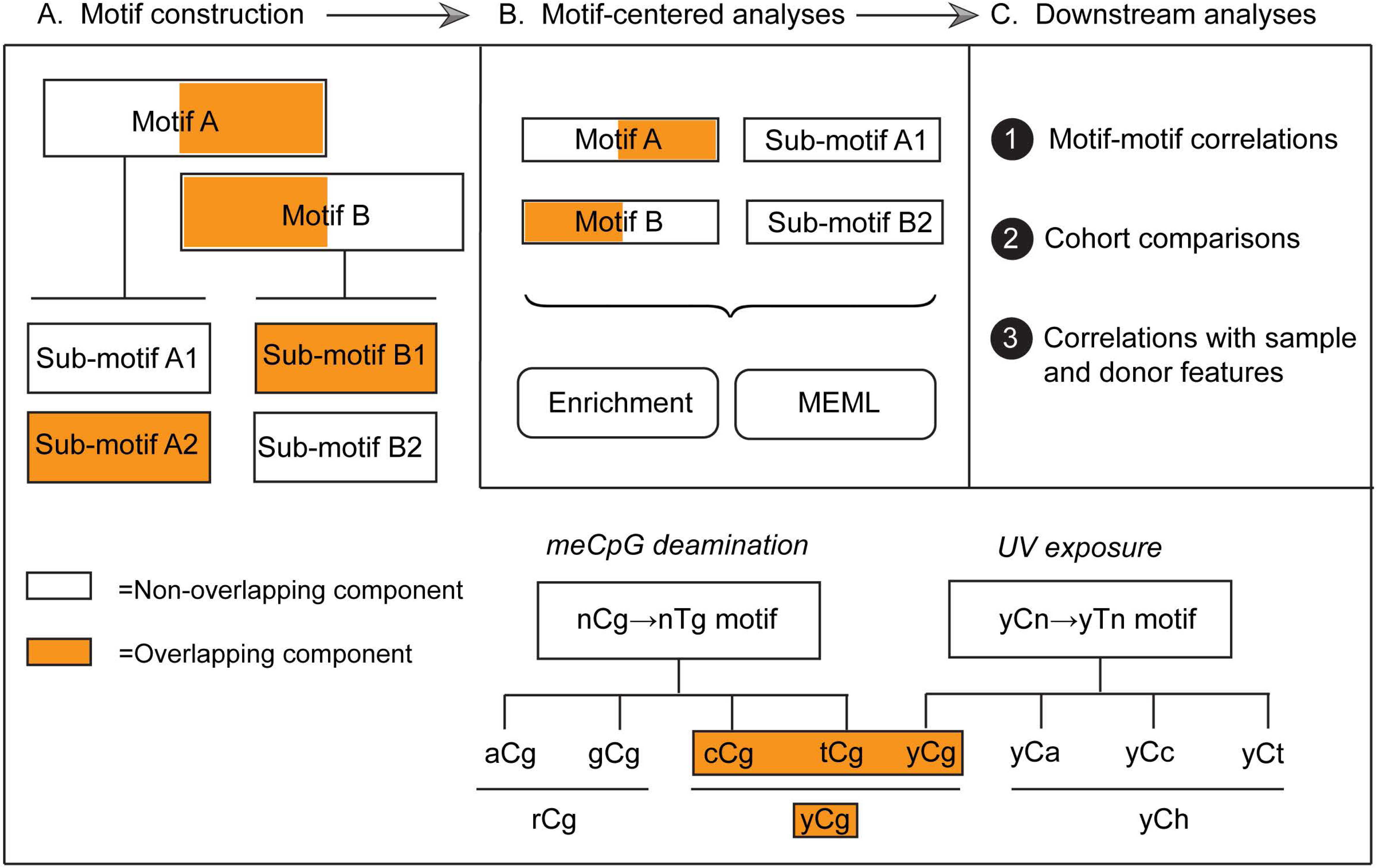
Framework of analyses performed in this study. **A.** Motifs and sub-motifs are constructed based on mechanistic knowledge about the trinucleotide preference of different mutagenic processes, as shown for dissection of motifs associated with meCpG deamination and UV exposure. **B.** Motif-centered analyses are performed on the mutation catalogue of each sample to detect enrichment and MEML attributable to the associated mutagenic process. **C.** Enrichment and MEML values are utilized in downstream analyses to find correlations with other motifs, sub-motifs, and with the features of individual samples and donors, and to perform comparisons between different cohorts.

### Motif-centered analyses can identify mutagenic mechanisms operating universally or in a tissue-specific manner

The most ubiquitous motif identified in multiple samples across all tissue types was nCg→nTg associated with cytosine deamination in 5meCpG (Fig. 2 and Supplementary Fig. S2). The four trinucleotides built into the nCg→nTg mutational motif (C→T mutations in aCg, cCg, gCg, and tCg) mimic the main peaks in COSMIC signature SBS1, a signature reliably extracted by different agnostic signature extraction algorithms in somatic and cancerous tissue types. Both SBS1 signature and nCg→nTg motif are age dependent, conforming to the expectation for endogenous mutational process of non-enzymatic deamination and providing additional assurance in their mechanistic assignment.

Another ubiquitous motif revealed by our analyses in healthy and diseased normal tissues was aTn→aCn associated with mutagenesis by epoxides and other S_N_2 reacting electrophiles (Fig 2), which was also identified in virtually all cancer types (41). Analyses utilizing sub-motif construction and motif-motif correlation further separated aTn mutagenesis from confounding input of T→C mutations by UV-mutagenesis (Fig. 4C). Peaks of aTn motif closely matched the peaks of COSMIC signature SBS16 extracted from large PCAWG dataset of human cancer mutations (Supplementary Fig. S3A). However, SBS16 does not have an assigned etiology and was extracted from only two cancer types (head and neck and liver) (18). Several studies extracted SBS16 from a limited number of somatic tissue types. SBS16 was found in neurons (46, 71), cirrhotic liver (72), esophagus (17), lung (73), and breast (74). Of note, SBS16 was extracted from breast tissues by only one out of three different signature extraction algorithms utilized, and was reported as a low-confidence signature by the authors (74). SBS16 was also only detected in 3 out 24 healthy tissues and 3 out of 14 diseased tissues by refitting of somatic mutations analyzed in this study with the published COSMIC signatures (Fig 1, Supplementary Fig. S1 and S2). Sample-specific signature refitting detected SBS16 in most tissues but in a much smaller number of samples than the aTn motif analysis (Supplementary Fig. S4). These observations suggest two possibilities worth further experimental investigation: either SBS16 captures the same or a subset of the same mutagenic activity as aTn motif but is detected less efficiently by signature extraction methods due to their known limitations, or the correlation between aTn and SBS16 reflects shared trinucleotide sequence similarity rather than a common mechanistic origin.

Framework of motif centered analysis further proved to be productive in identifying tissue-specific mutation mechanisms, as exemplified by the detection of UV and APOBEC mutagenesis. UV-associated mutational motifs yCn and nTt were used based on prior mechanistic evidence and were found only in skin (Fig. 8). Sub-motif analyses allowed to rule out UV-like mutagenesis as confounded by other mechanisms in several other tissues where yCn motif was detected.

Peaks corresponding to trinucleotide motifs yCn and nTt are detectable among the 96-motif spectra of UV-associated COSMIC signatures SBS7a,b and SBS7c,d, respectively, however other peaks were also present in these signatures (Supplementary Fig. S3B). Trinucleotides in APOBEC-associated motif tCw→tTw corresponds to SBS2, while tCw→tGw corresponds to SBS13 peaks (Supplementary Fig. S3B). Motif-centered analyses identified APOBEC mutagenesis in bladder, liver, lung, breast, and small intestine (Fig. 9, Supplementary Fig. S12), as opposed to signature refitting which identified low contribution of SBS2 and SBS13 only in bladder. (Fig 1B, Supplementary table S3c). Construction of tetranucleotide motifs based on prior knowledge about motif preference by APOBEC3A and APOBEC3B (36) allowed the assignment of APOBEC3A as the predominant mutator in tissues where APOBEC mutagenesis was identified (Supplementary Fig. S12), an outcome not achievable by signature analyses.

We also detected sporadic presence of some motifs in multiple tissues and diseases. Samples with enrichment of gCn→gAn motif, which is associated with acetaldehyde exposure and found primarily enriched in liver cancers (59), were detected in multiple somatic tissue types analyzed in our study (12 out of 24; Supplementary Fig. S11A, Supplementary table S9). cTg→cAg motif associated with exposure to aristolochic acid (AA) was detected in healthy colon and bladder, and in both healthy and diseased samples of liver and lung (Supplementary Fig. S11E, Supplementary table S9). SBS22a and SBS22b are associated with AA exposure and contain peaks mimicking cTg→cAg motif among other peaks. SBS22 has been extracted from cancers in upper urinary tract, liver, kidney, and from lung cancer in never-smoking East Asian individuals (75, 76). SBS22 was also extracted from three healthy liver tissue donors and from donors with alcohol-related liver diseases (61, 72). To our knowledge, this is the first indication of AA-induced mutagenesis in normal lung tissues which was further exacerbated in donors with COPD.

We note that the estimated motif-associated MEML can be reduced by the number of mutational mechanisms operating and preferentially mutagenizing the same base in a sample. However, calculation of the minimum estimate of mutation load in a motif provides a baseline value associated with a mutagenic process that can be compared against the baseline value in another sample. Further limitations of motif-centered analyses are due to the restrictive availability of knowledge about high confidence, experimentally validated motifs and the unavoidable overlap between some motifs that cannot be resolved by sub-motif approach. Signature extraction method on the other hand has unique capability to find new mutation patterns amenable for hypothesis generation. Signature extraction solves an optimization problem, while motif-centered analysis performs statistically stringent hypothesis testing against mechanistically defined expectations. We therefore view both approaches as complementary and essential in identifying and validating sources of somatic mutations. Efforts from the broader research community is needed to discover and characterize novel, mechanistically grounded motifs that may complement signatures with and without known etiologies and expand the repertoire of mutagenic processes that can be reliably detected and quantified.

### Motif-specific mutational burdens can serve as indicators of an individual’s inherited susceptibilities and cumulative exposures over the lifespan

We explored all motifs and all datasets for potential association between somatic mutagenesis and disease, and found several diseases associated with higher prevalence of motif-specific mutation burdens (Table 2).

**Table 2.** Summary of somatic mutagenesis in non-cancer diseases compared to their corresponding healthy tissues detected by motif-centered analyses.

| Tissue | Disease name | Disease specific physiology | aTn<br>Small epoxides,<br>S<sub>N</sub>2 electrophiles | nCg<br>meCpG<br>deamination | hTg<br>MMS, S<sub>N</sub>2<br>electrophiles | gCn<br>Acetaldehyde | cTg<br>Aristolochic<br>acid | tgC<br>Redox |
| --- | --- | --- | --- | --- | --- | --- | --- | --- |
| Brain | Alzheimer's disease | Amyloid plaques and tau tangles (not fully understood) | Decreased mutations<sup>1</sup> | Decreased mutations<sup>1</sup> | - | - | - | - |
| Brain | Tourette's Syndrome | Neurotransmitter disruption (not fully understood) | - | Clock-like<sup>2</sup><br>- | - | - | - | - |
| Colon | Crohn's disease | Chronic inflammation; autoimmune | Clock-like<sup>2</sup><br>- | - | - | - | - | - |
| Colon | Ulcerative colitis | Chronic inflammation; autoimmune | Clock-like<sup>2</sup><br>- | Clock-like<sup>2</sup><br>Increased mutations<sup>3</sup> | - | Increased mutations | - | - |
| Lung | Chronic Obstructive Pulmonary Disease | Chronic inflammation | Increased mutations | Increased mutations | Increased mutations | Increased mutations | Increased mutations | - |
| Liver | Alcoholic Cirrhosis | Chronic inflammation, fibrosis | Increased mutations | Increased mutations | - | - | Increased mutations<sup>4</sup> | - |
| Liver | Primary Sclerosing Cholangitis | Chronic inflammation; autoimmune | Increased mutations<sup>4</sup> | Increased mutations<sup>4</sup> | - | - | - | - |
| Liver | Non-Alcoholic Steatohepatitis | Chronic inflammation, fat accumulation | Clock-like<sup>2</sup><br>Increased mutations | Clock-like<sup>2</sup><br>Increased mutations | - | - | - | - |
| Small intestine | Coeliac | Autoimmune reaction to gluten | - | - | - | - | - | Increased mutations |

<sup>1</sup> In upper age tercile of one study
<sup>2</sup> In the diseased cohort of the specified tissue
<sup>3</sup> In lower age tercile
<sup>4</sup> Significant before FDR correction
- No difference between healthy and diseased donors

Importantly, increased mutagenesis was found in only some of all diseases of a specific tissue type, suggesting an involvement of disease-specific physiological causes as opposed to a strictly age-dependent increase in mutagenesis. Several diseases associated with motif-specific mutagenesis are also associated with increased inflammation. The excess of mutations in non-neoplastic inflammatory diseases can stem from a higher rate of exposure and DNA damage compared to healthy tissues. Continued exposure to xenobiotic sources of DNA damage, like smoking in COPD patients and alcohol consumption in cirrhosis patients, increases the load of mutagenic lesions. Xenobiotic precursors can be endogenously metabolized into reactive small epoxides by CYP2E1 and other CYP epoxygenases primarily in the liver and released back into the body (77, 78). Reactive oxygen and nitrogen species (RONs) produced in inflammatory diseases can overwhelm the DNA damage repair pathways and interfere with DNA repair efficiency (79). Reactive oxygen species may further promote deamination of meCpG dinucleotides (80).

The lower motif-associated mutation loads detected in brain samples of donors with Alzheimer’s disease compared to healthy donors could be explained by *in situ* selection against highly mutated neurons and preferential sequencing of comparatively fit, lower burden neurons. This trend was observed in another post-mitotic cell type, articular chondrocytes, where samples from healthy donors showed higher or comparable total mutation and motif-associated mutation loads compared to samples from donors with osteoarthritis (81, 82). Decline in detection of motif-associated mutagenesis in post-mitotic cells may be indicative of their limited access to DNA damage repair machinery.

The effect of age and disease on somatic mutagenesis in normal tissues can also be supplemented by the impact of individuals’ genotype, which is well-established in cancers (83) and has also been shown for several normal tissues (84, 85). Specifically, impact of germline genotype was established on mutagenic processes in cancers that are associated with somatic mutagenesis motifs nCg (meCpG deamination) and tCw (APOBEC) revealed in this study. Germline and somatic variants in mismatch repair complex MutSα are associated with hypermutation of meCpG motifs in cancers (86). Germline mutant alleles of meCpG binding glycosylase MBD4 also predispose to meCpG mutagenesis in cancers (32, 87, 88). Common polymorphisms in APOBEC3 gene cluster cause predisposition to cancer and exacerbate APOBEC mutation load in tumors (26, 32, 89, 90). Thus, motif-associated mutations can be utilized as a phenotype for germline association studies to identify predisposition and vulnerability to specific mutagenic processes.

Since motif-specific somatic mutagenesis are mechanistically linked to their sources, it allows quantification of the activities of endogenous and environmental factors that increase somatic mutation burden. Detection and monitoring of these mutagenic activities in normal tissues can prompt early diagnosis of cancer and non-cancerous diseases, which in turn may facilitate disease prevention, management, and improved treatment outcomes.

## Data access

All mutation calls analyzed in this study are available to download in VCF format at https://vijglab.einsteinmed.edu/SomaMutDB/download/ or in the supplementary material of the indicated papers. All data generated in this study, including numerical values underlying figures and the summary statistics are in the Supplementary Tables. P-MACD output for APOBEC cluster analyses are available in Supplementary Data S1. P-MACD package to perform motif-centered analyses is deposited at https://doi.org/10.5281/zenodo.8018096; all versions of P-MACD and future updates are available at https://github.com/NIEHS/P-MACD.

## Acknowledgements

This work was supported by the NIH, National Institute of Environmental Health Sciences Intramural Research Program Project Z1AES103266 to D.A.G. The contributions of the NIH authors are considered Works of the United States Government. The findings and conclusions presented in this paper are those of the authors and do not necessarily reflect the views of the NIH or the U.S. Department of Health and Human Services. We are thankful to Dr. Ashley Brooks for help with vcf2maf tool, to Dr. Min Shi for consultation on multivariable regression analyses, to Drs Rajula Alleva-Elango and Natalya Degtyareva for advice on manuscript, and to Lois Wyrick for help with illustrations.

## Author contributions

S.M.S-Conceptualization (Lead), Data curation (Lead), Formal analysis (Lead), Investigation (Equal), Methodology (Equal), Software (Supporting), Visualization (Lead), Writing - original draft (Lead), Writing - review & editing (Lead)

Y.-C.H. – Formal analysis (Supporting), Software (Supporting)

L.J.K. – Software (Lead)

D.A.G. – Conceptualization (Lead), Formal analysis (Supporting), Funding acquisition (Lead), Investigation (Equal), Methodology (Equal), Project administration (Lead), Supervision (Lead), Writing - original draft (Equal), Writing - review & editing (Equal)

## Supplementary information

### Supplementary figures

**Supplementary Figure S1.**
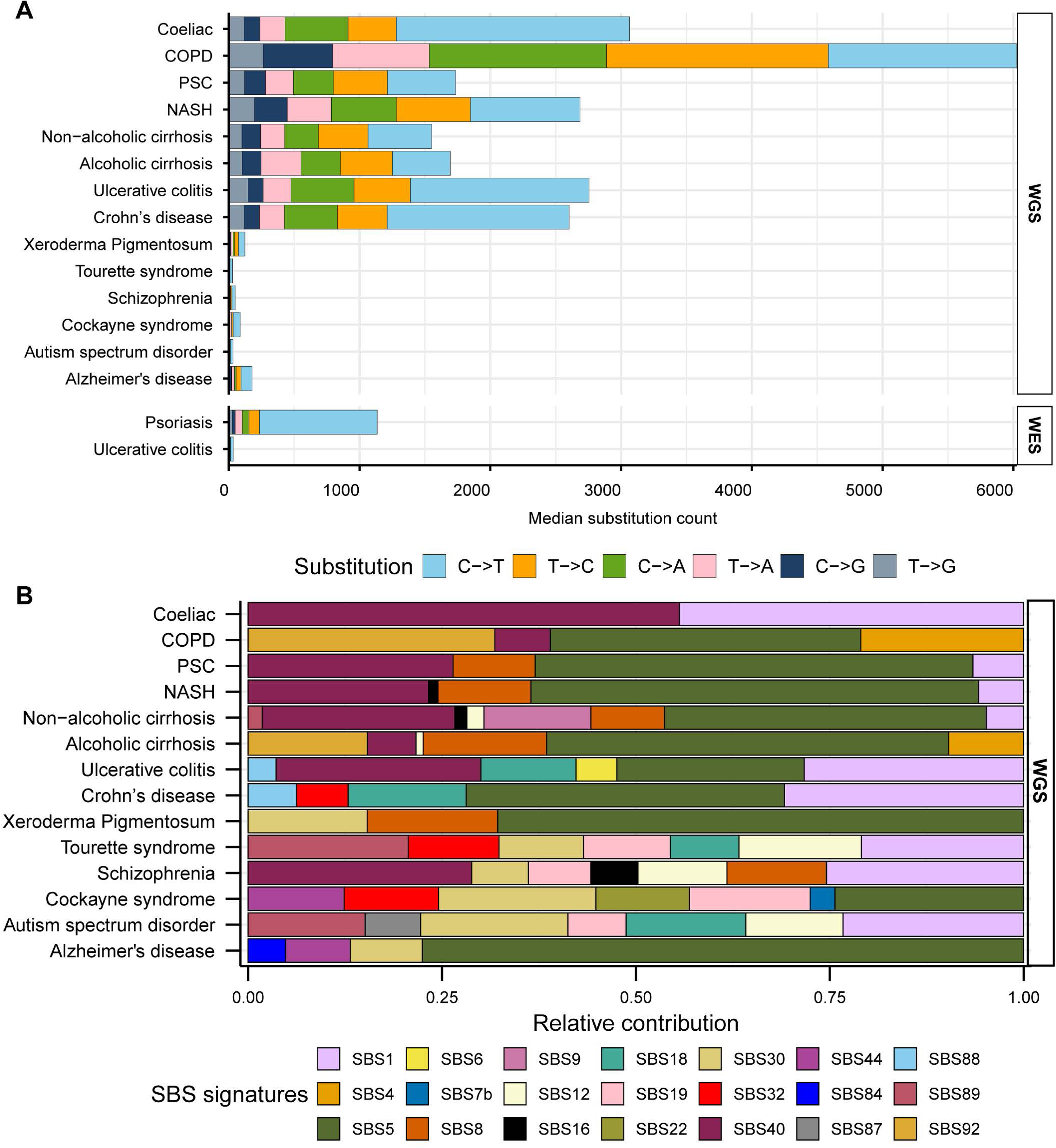
Somatic mutation spectra and contribution of COSMIC signatures in diseased tissues. **A**. Median counts for each base substitution including reverse complements are shown in descending order for diseased WGS and WES samples. **B.** Relative contributions of published COSMIC signatures in the mutation profiles of WGS diseased tissues. All source data are available in Supplementary Table S3.

**Supplementary Figure S2.**
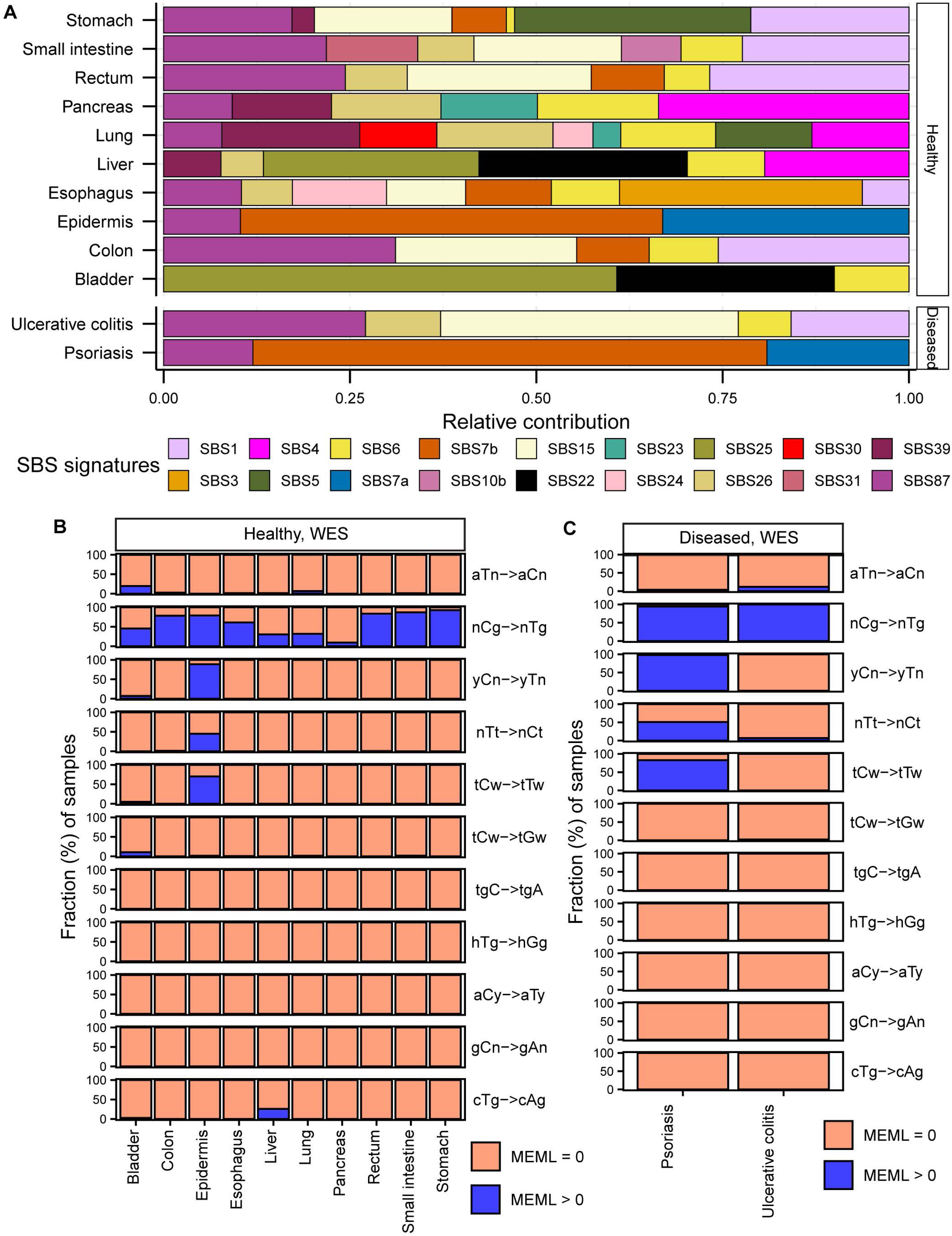
SBS signatures and mutational motifs in exome-sequenced samples. **A.** Relative contributions of published COSMIC signatures in the mutation profiles of WES samples are shown for healthy and diseased tissues. **B-C**. The percentages of samples with zero and non-zero MEML counts for the 11 known mutational motifs calculated for WES samples in healthy **(B)** and diseased **(C)** tissues. Source data are available in Supplementary Tables S3 and S4.

**Supplementary Figure S3.**
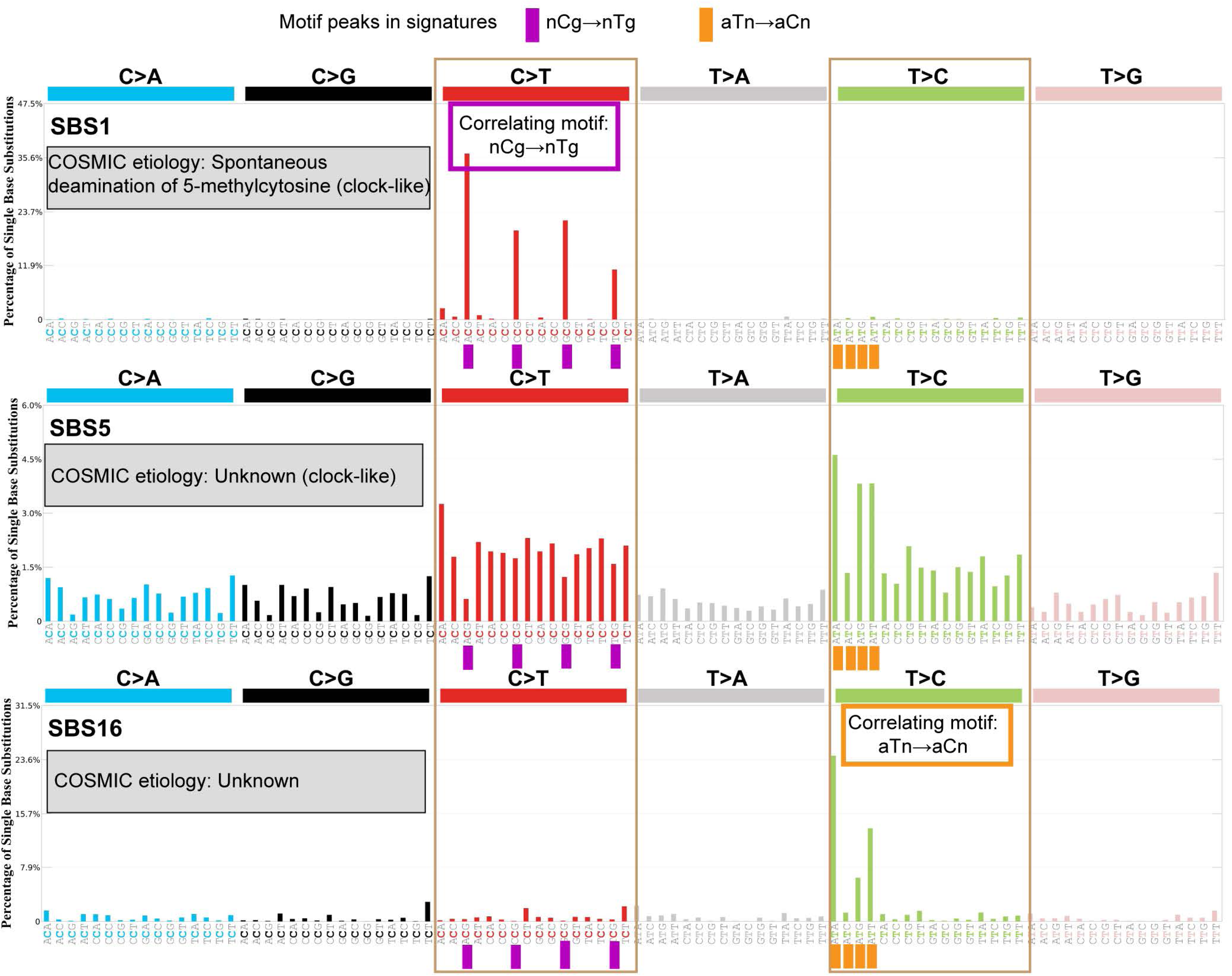
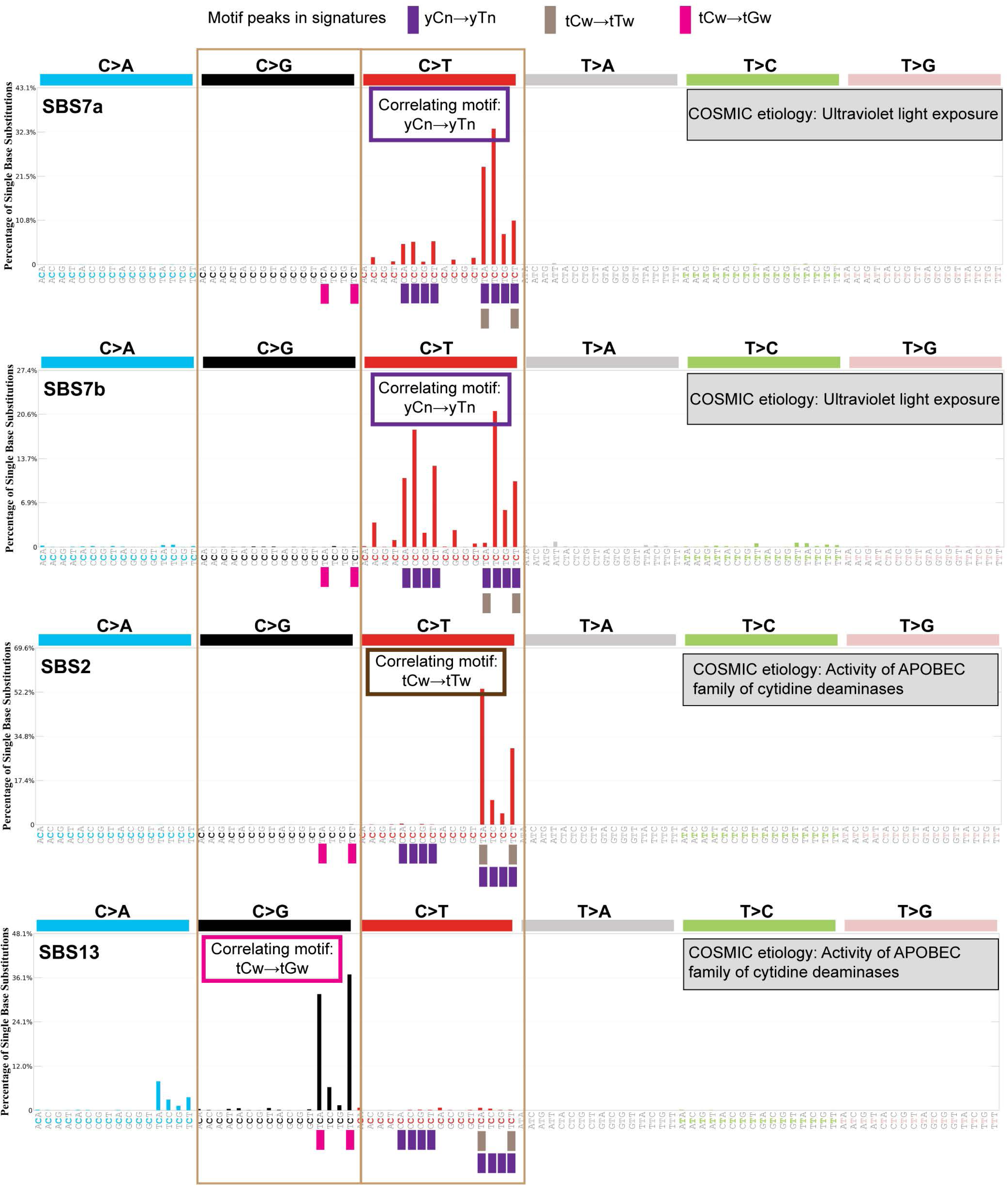
COSMIC signature profiles and correlating motifs. Plots displaying mutation profiles for GRCh38 with assigned etiologies were downloaded from COSMIC website and shown for signatures **(A)** SBS1, SBS5, SBS16, **(B)** SBS7a, SBS7b, SBS2, and SBS13. Motifs correlating with the signatures are indicated on each plot with vertical bars under corresponding motif peaks in signature spectra. Bar colors indicate individual motifs.

**Supplementary Figure S4.**
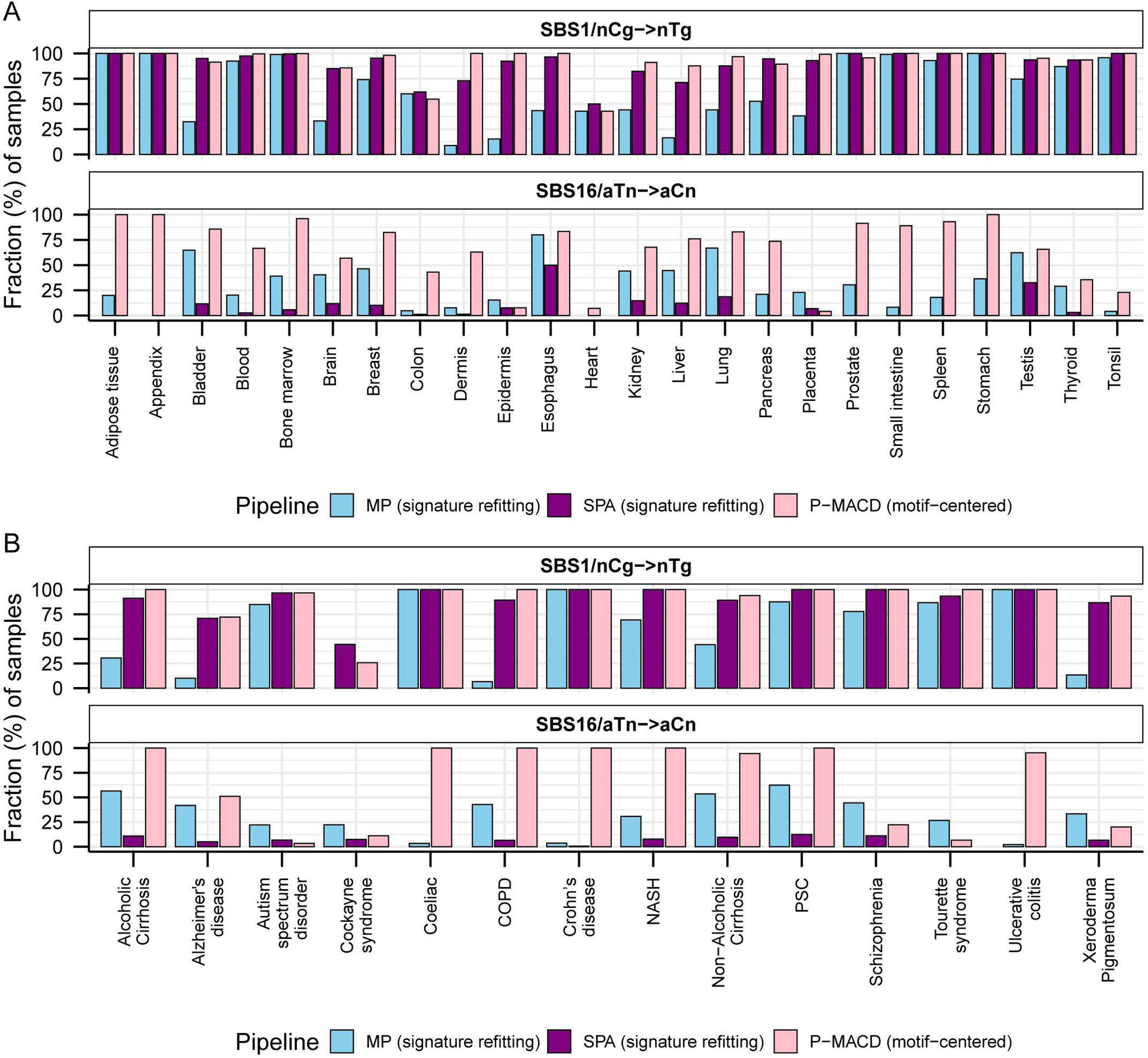
Sample-specific detection sensitivity of signature refitting tools and motif-centered analysis. Percentage of samples with detectable SBS1 signature/nCg motif and SBS16 signature/aTn motif in **(A)** healthy and **(B)** diseased tissues are shown using two signature refitting tools and P-MACD, the tool for motif-centered analyses. Source data are available in Supplementary Table S4.

**Supplementary Figure S5.**
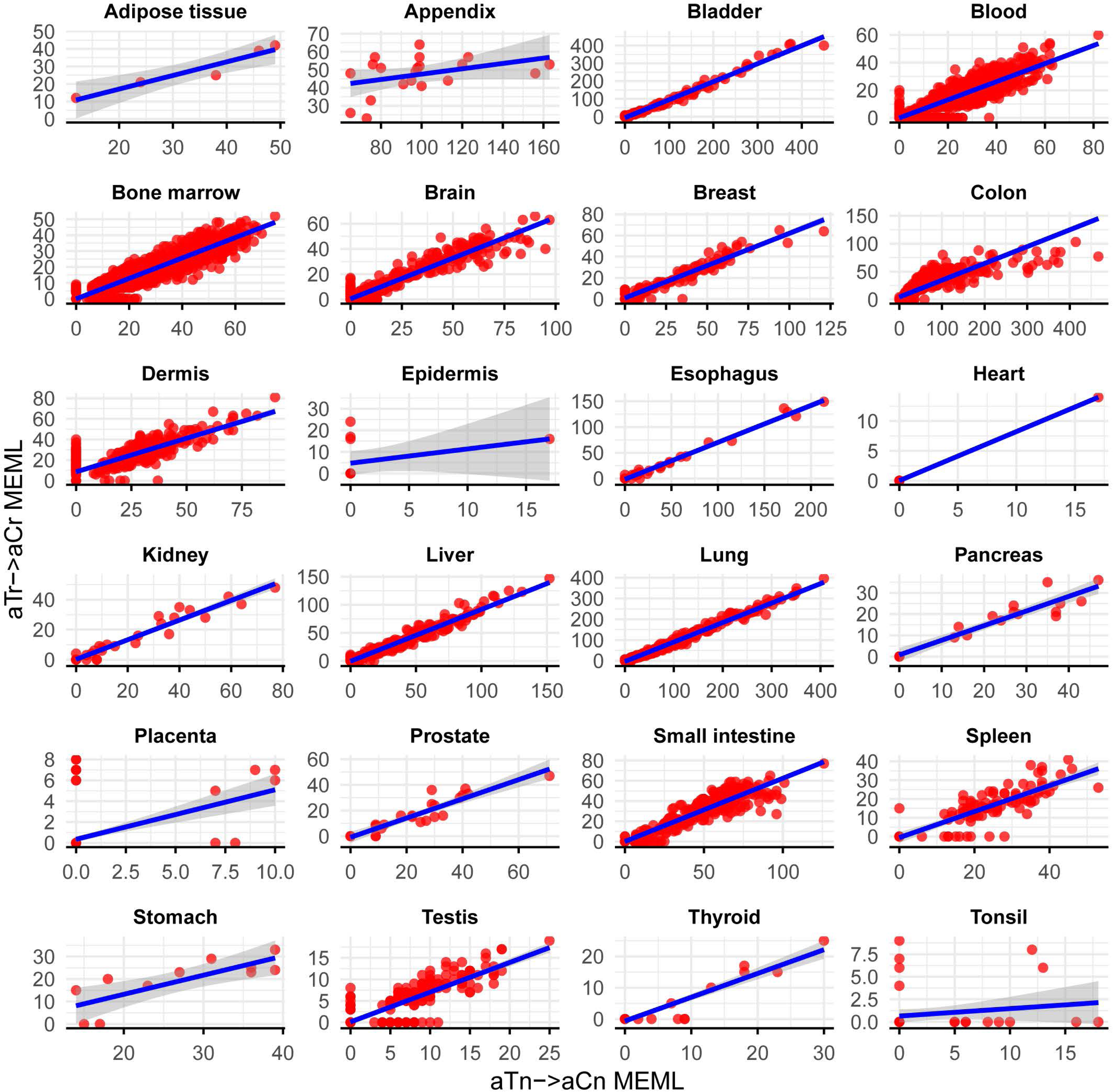
Analysis of aTn→aCn sub-motifs to detect true presence of aTn motif in healthy tissues. Scatter plots showing aTn motif MEML plotted against aTr motif MEML in WGS samples across different tissues. Blue line indicates line of best fit from a simple linear model.

**Supplementary Figure S6.**
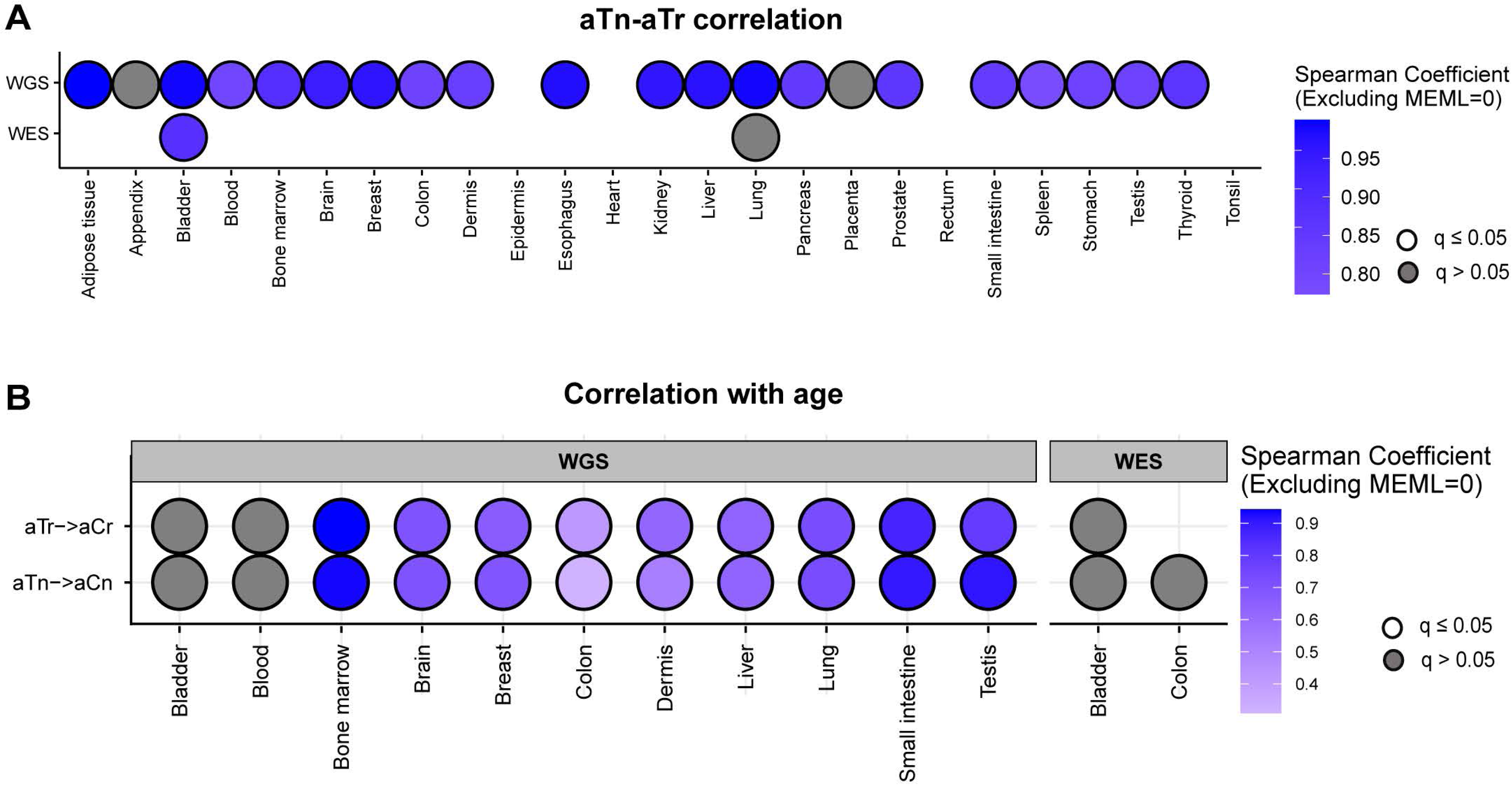
Analyses of aTn motif and sub-motifs in samples with non-zero MEML values. **A.** Correlation matrix showing Spearman correlation between aTn→aCn MEML and aTr→aCr MEML excluding either MEML =0 to identify true aTn mutational motif. Grey circles indicate P value >0.05 after correction for multiple hypothesis using Benjamini-Hochberg method within each sequencing group (WGS or WES). No circles indicate insufficient samples with MEML>0 for a tissue type to perform correlation analyses. **B.** Correlation matrix showing donor age versus mean donor MEML of aTn and aTr motifs excluding either MEML=0. Spearman correlations were performed between donor age and mean MEML count of all samples from an individual donor for the indicated tissues and mutational motifs. Only tissues with sufficient number of donors to support age-correlation analyses are shown. Grey circles indicate P value >0.05 after correction for multiple hypothesis using Benjamini-Hochberg method for a given motif within each sequencing group (WGS or WES). No circles indicate insufficient individual donors with mean MEML>0 for to perform correlation analyses. All source data and statistical analyses can be found in Supplementary Table S5.

**Supplementary Figure S7.**
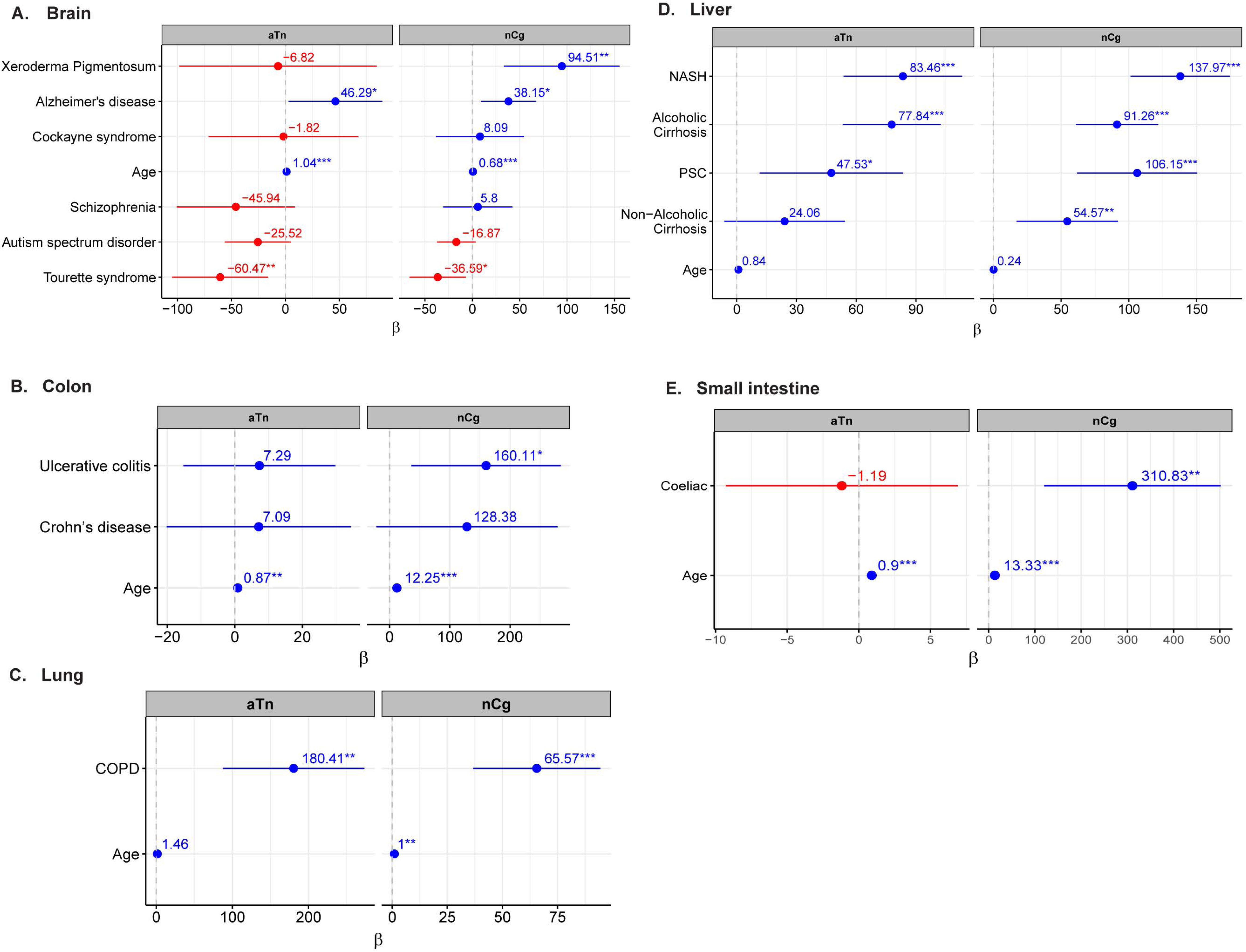
Multivariable regression analyses of aTn and nCg motifs in non-cancer diseases. *β*-coefficients from multivariable regression analyses using donor age and disease status as predictor variables are plotted for brain **(A)**, liver **(B)**, colon **(C)**, lung **(D)**, and small intestine **(E)**. *P value ≤ 0.05, **P value ≤ 0.01, ***P value ≤ 0.001. Source data are available in Supplementary Tables S6c and S8d.

**Supplementary Figure S8.**
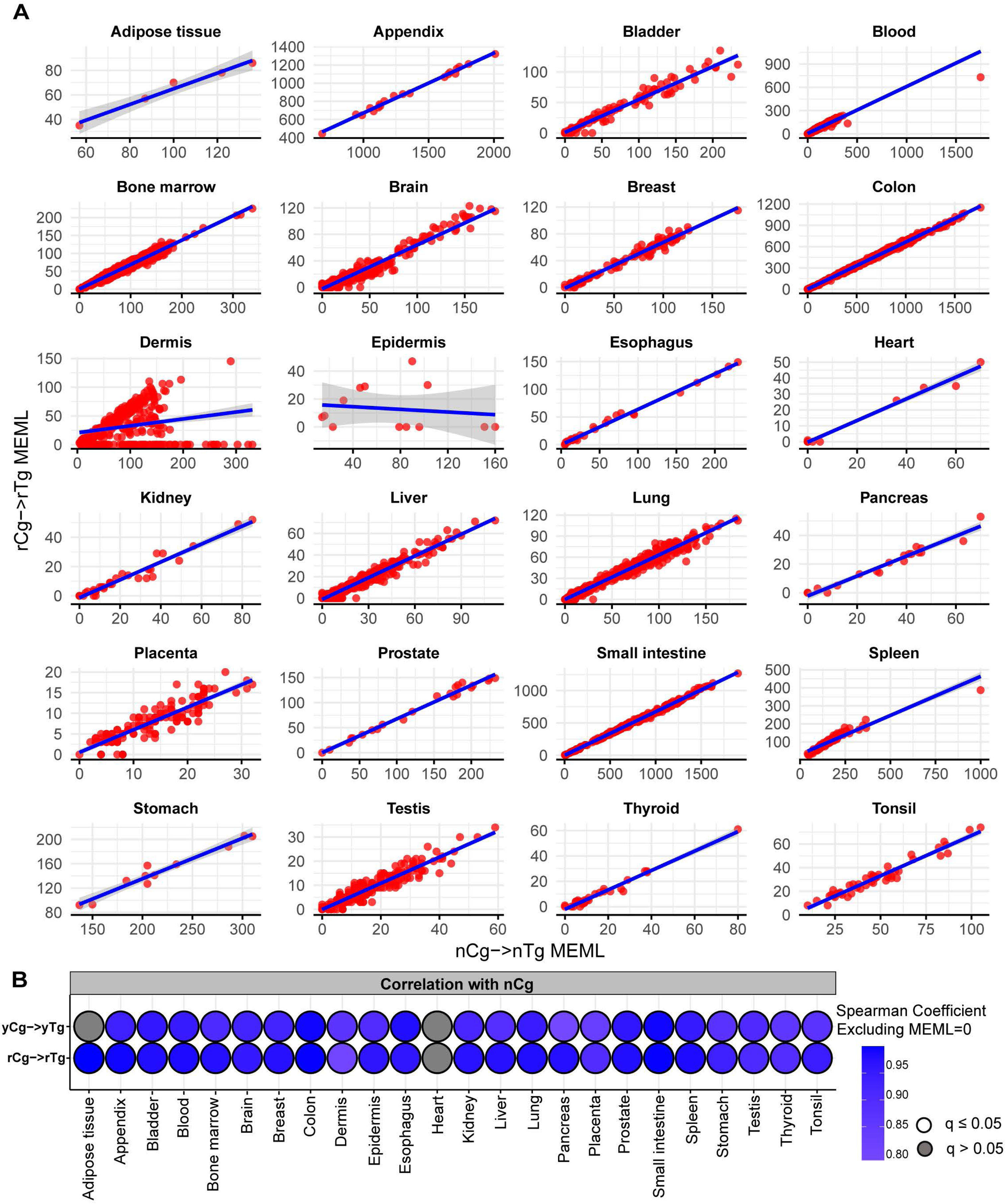
Analysis of nCg→nTg sub-motifs and their correlation with each other in healthy tissues. **A.** Scatter plot showing nCg→nTg MEML plotted against MEML of sub-motif rCg→rTg, the component of nCg non-overlapping to UV-motif yCn, in WGS samples across different tissues. Blue line indicates line of best fit from a simple linear model. **B.** Correlation matrix showing Spearman correlation of nCg→nTg with sub-motifs rCg→rTg (UV non-overlapping) and yCg→yTg (UV overlapping), excluding either MEML =0. Grey circles indicate P value >0.05 after correction for multiple hypothesis using Benjamini-Hochberg method within each sequencing group (WGS or WES). No circles indicate insufficient samples with MEML>0 for a tissue type to perform correlation analyses. C. Scatter plot showing nCg→nTg MEML plotted against MEML of yCh→yTh, the component of UV-motif yCn non-overlapping to nCg, in WGS samples of dermis and epidermis. Blue line indicates line of best fit from a simple linear model. All source data and statistical analyses can be found in Supplementary Table S7.

**Supplementary Figure S9.**
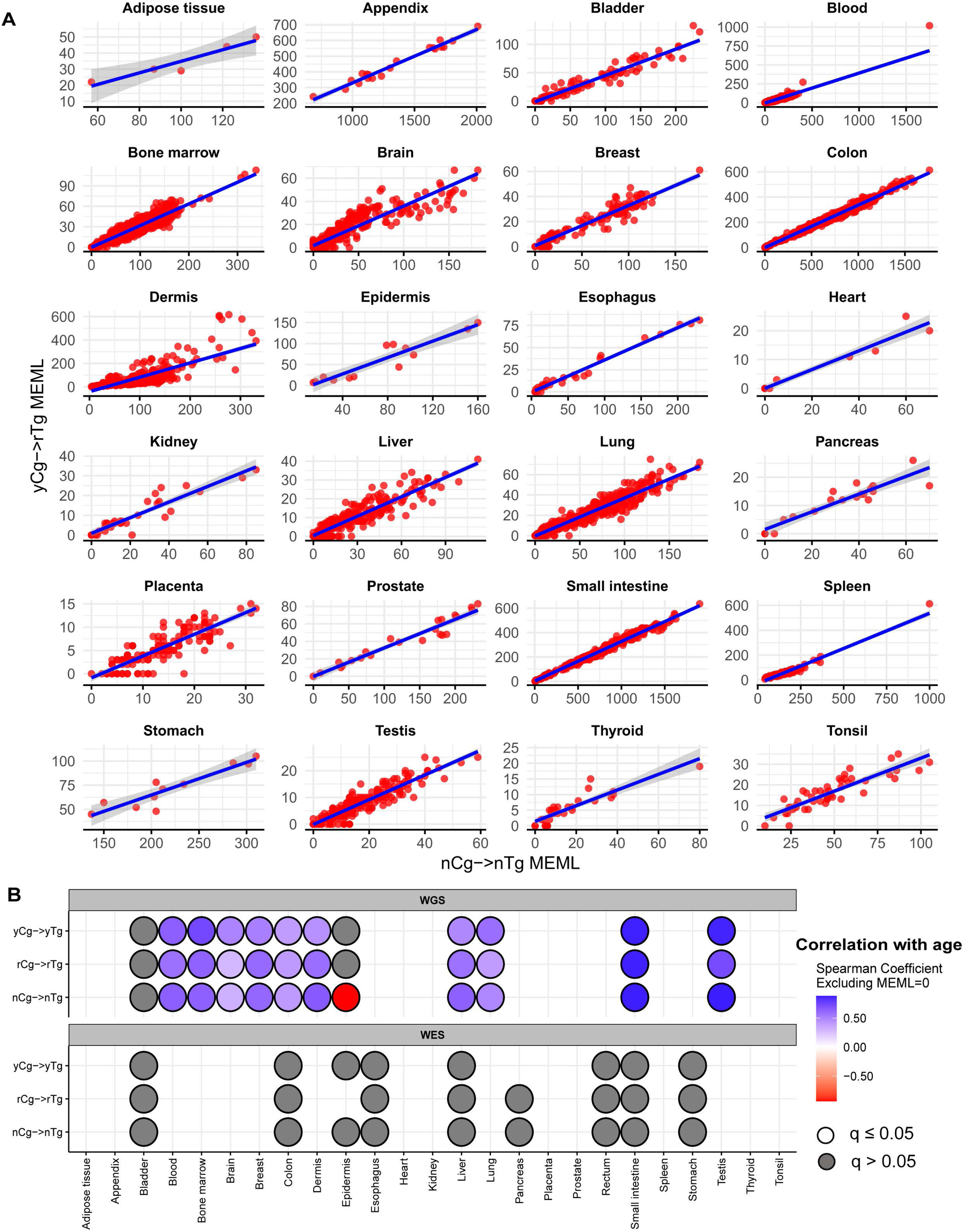
Analysis of nCg→nTg sub-motifs and their correlations with donor age in healthy tissues. **A.** Scatter plot showing nCg→nTg MEML plotted against MEML of sub-motif yCg→yTg, the component of nCg overlapping to UV-motif yCn, in WGS samples across different tissues. Blue line indicates line of best fit from a simple linear model. **B.** Correlation matrix showing Spearman correlation of donor age with donor-specific mean MEML value for nCg→nTg and sub-motifs rCg→rTg (UV non-overlapping) and yCg→yTg (UV overlapping), excluding either MEML =0. Grey circles indicate P value >0.05 after correction for multiple hypothesis using Benjamini-Hochberg method within each sequencing group (WGS or WES). No circles indicate insufficient samples with MEML>0 for a tissue type to perform correlation analyses. All source data and statistical analyses can be found in Supplementary Table S7.

**Supplementary Figure S10.**
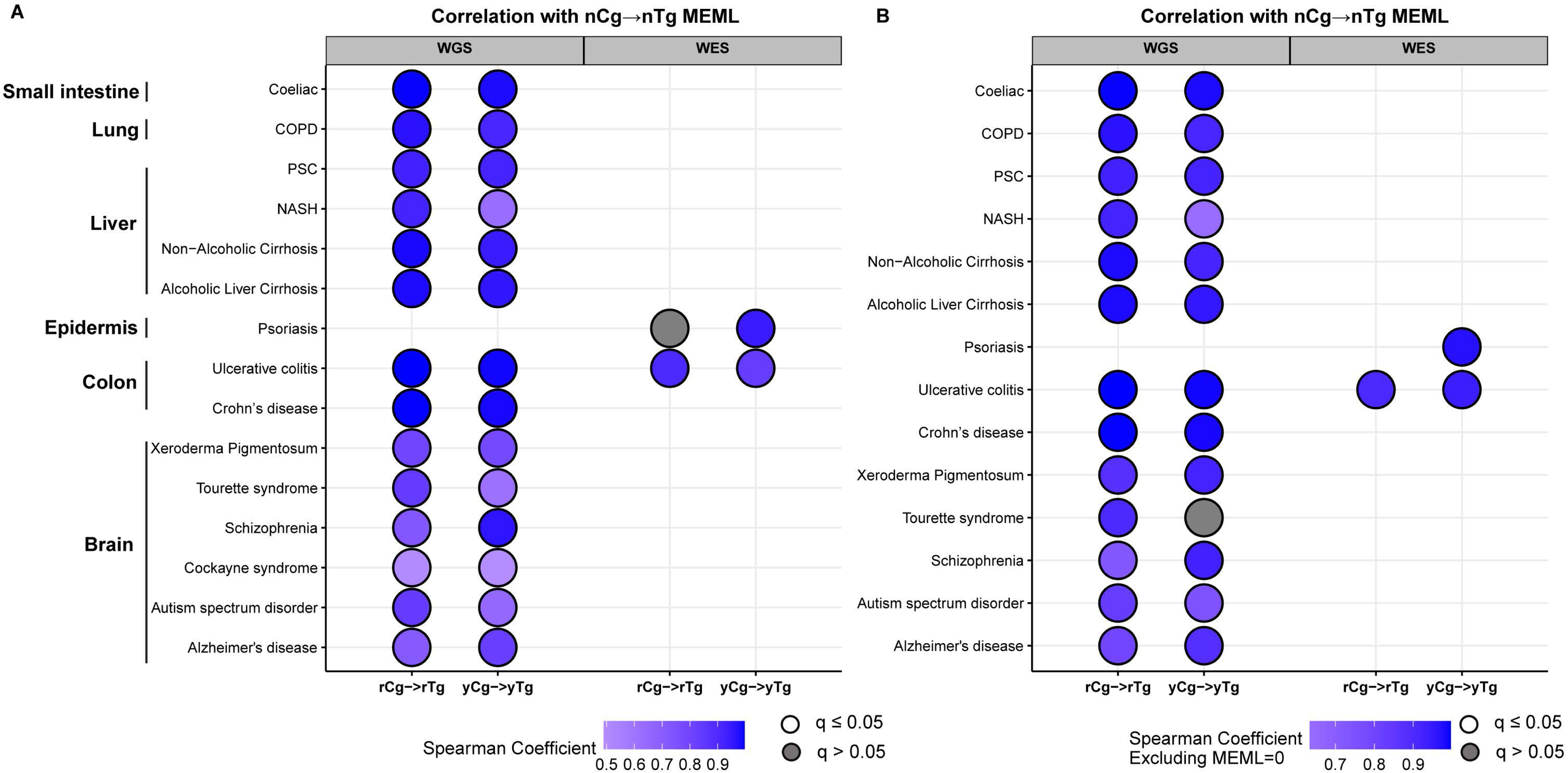
Analyses of sub-motifs to detect true spontaneous meCpG cytosine deamination-mediated nCg→nTg in diseased tissues. Correlation matrix showing Spearman correlation of nCg→nTg with sub-motifs rCg→rTg (UV non-overlapping) and yCg→yTg (UV overlapping), including MEML = 0 **(A)** and excluding pairs with either member’s MEML =0 **(B)** in different disease categories. Grey circles indicate P value >0.05 after correction for multiple hypothesis using Benjamini-Hochberg method within each sequencing group (WGS or WES). No circles indicate insufficient samples with MEML>0 for a tissue type to perform correlation analyses. Source data are available in Supplementary Table S8b.

**Supplementary Figure S11.**
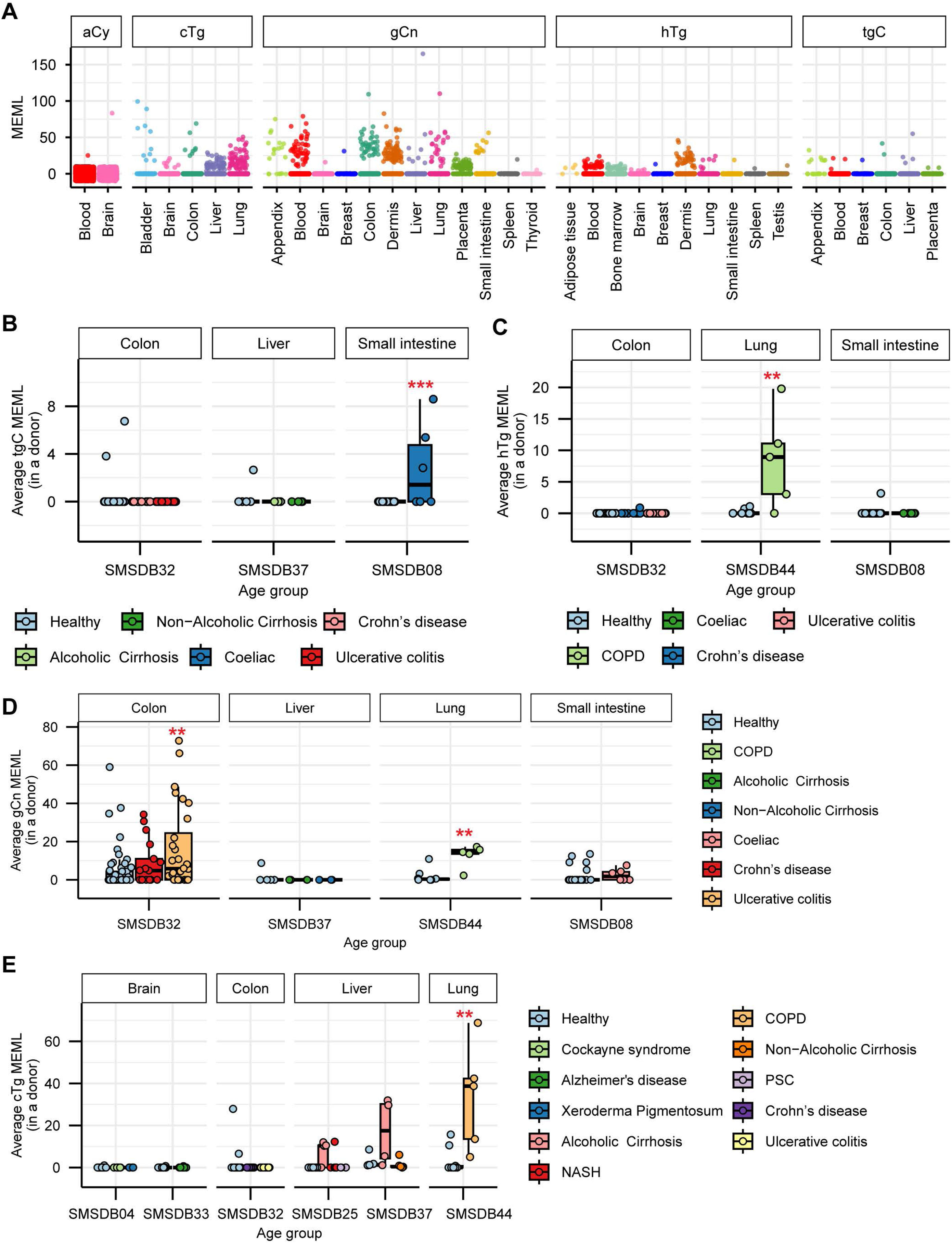
Analysis of sporadic motifs in healthy and diseased tissues. **A.** Sample-specific MEML counts shown in jitter plots for motifs indicated in panel labels for WGS samples of healthy tissue types. **B-E.** Mean donor MEML for healthy and diseased samples are shown in a boxplot for tgC (B), hTg (C), gCn (D) and cTg (E) motifs. X-axes indicate source study, and panel labels indicate tissue type of the samples. Analyses of only WGS samples are shown. Asterisks in boxplots represent statistical significance (Wilcoxon Rank Sum test) compared to the healthy donors within each study after correcting for multiple hypothesis testing using Benjamini-Hochberg method. *P value ≤ 0.05, **P value ≤ 0.01, ***P value ≤ 0.001. All source data and statistical analyses can be found in Supplementary Table S9.

**Supplementary Figure S12:**
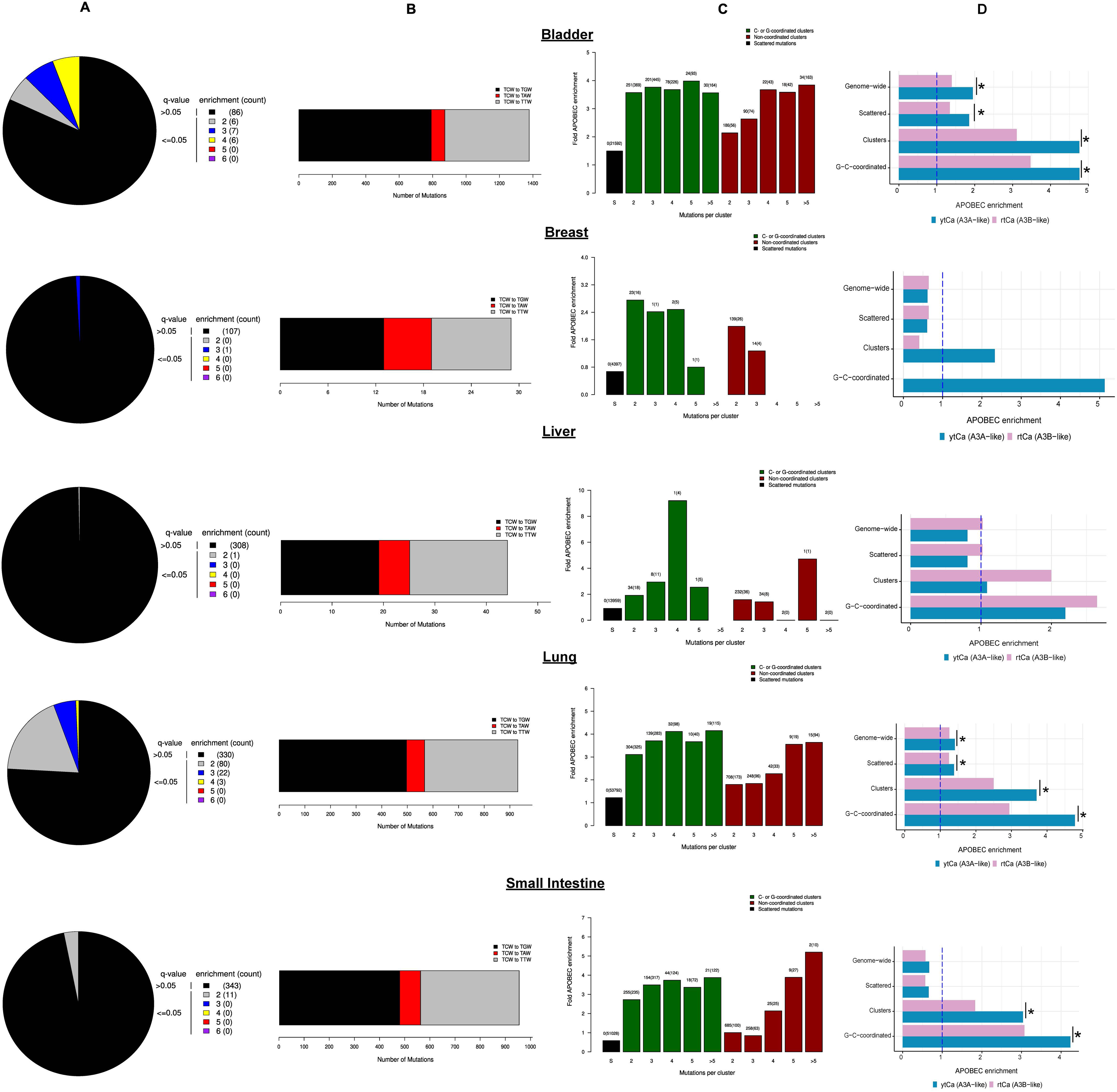
Tissues with multiple features indicating APOBEC mutagenesis. **A.** Fold-enrichment of APOBEC mutagenesis motif over the expected occurrence of random mutagenesis in the indicated tissues. Samples are categorized into color coded bins based on their FDR corrected q-values (Benjamini-Hochberg) and fold enrichment values. Fold enrichment bin sizes are increments of 1 unit. All samples displaying a q-value >0.05 are placed in one bin (black) regardless of the fold APOBEC enrichment in the sample. The maximum fold enrichment for each bin is indicated in the figure legend, with the number of samples in its category shown in parentheses. **B.** The number of mutations categorized into three possible types of substitutions at the tCw motif (complementary mutations included) within C- or G-coordinated clusters in the indicated tissues. **C.** Fold enrichment of the APOBEC motif mutations in clusters of different sizes of two categories – C- or G-coordinated, non-coordinated, as well as in non-clustered (scattered) mutations in the indicated tissues. Values above the bars display the total number of mutation clusters in each class and in parentheses the total numbers of APOBEC motif mutations in a category. **D.** Enrichment of ytCa (APOBEC3A-like) motif and rtCa (APOBEC3B-like) motif among genome-wide, scattered, clustered, and G- or C-coordinated clustered mutations in the indicated tissues. The blue dashed vertical line indicates level of no enrichment. Where enrichment of both ytCa and rtCa motifs were statistically significant (Fisher’s Exact P-value ≤0.05), Breslow-Day test of homogeneity was performed to compare the odds ratios of the two motif enrichments. Asterisks indicate significant difference at P-value ≤0.05. Source data for plots **A**, **B**, and **C** are available in Supplementary Data S1. Source data and statistical analyses for ytCa and rtCa enrichment **(D)** can be found in Supplementary Table S11e and S11f.

### Supplementary Tables

**Supplementary Table S1:** Origins of mutation calls for samples analyzed in this study.

**Supplementary Table S2:** Sub-motifs of knowledge-based motifs with overlapping and non-overlapping components.

**Supplementary Table S3:** Somatic mutation spectra and COSMIC SBS signatures in normal cells. Contains numeric values underlying Figure 1, Supplementary Figures S1, S2a.

**Supplementary Table S4:** Prevalence of knowledge-based motifs in normal tissues. Contains numeric values underlying Figures 2, 3, Supplementary Figure S2b, S4.

**Supplementary Table S5:** aTn mutational motif in normal healthy cells. Contains numeric values underlying Figure 4, Supplementary Figures S5, S6.

**Supplementary Table S6:** Prevalence of aTn motif in diseased tissues. Contains numeric values underlying Figure 5, Supplementary Figure S7.

**Supplementary Table S7:** nCg mutational motif in normal healthy cells. Contains numeric values underlying Figure 6, Supplementary Figures S8, S9.

**Supplementary Table S8:** Prevalence of nCg mutational motif in diseased samples. Contains numeric values underlying Figure 7, Supplementary Figures S7, S10.

**Supplementary Table S9:** Prevalence of sporadic motifs in normal and diseased samples. Contains numeric values underlying Supplementary Figure S11.

**Supplementary Table S10:** UV mutational motif in normal cells. Contains numeric values underlying Figure 8.

**Supplementary Table S11:** APOBEC mutational motif in healthy normal tissues. Contains numeric values underlying Figure 9, Supplementary Figure S12.

### Supplementary Data

**Supplementary Data S1:** Raw output of P-MACD motif and cluster analyses to detect APOBEC mutagenesis. Contains numeric values underlying Supplementary Figure S12.

---

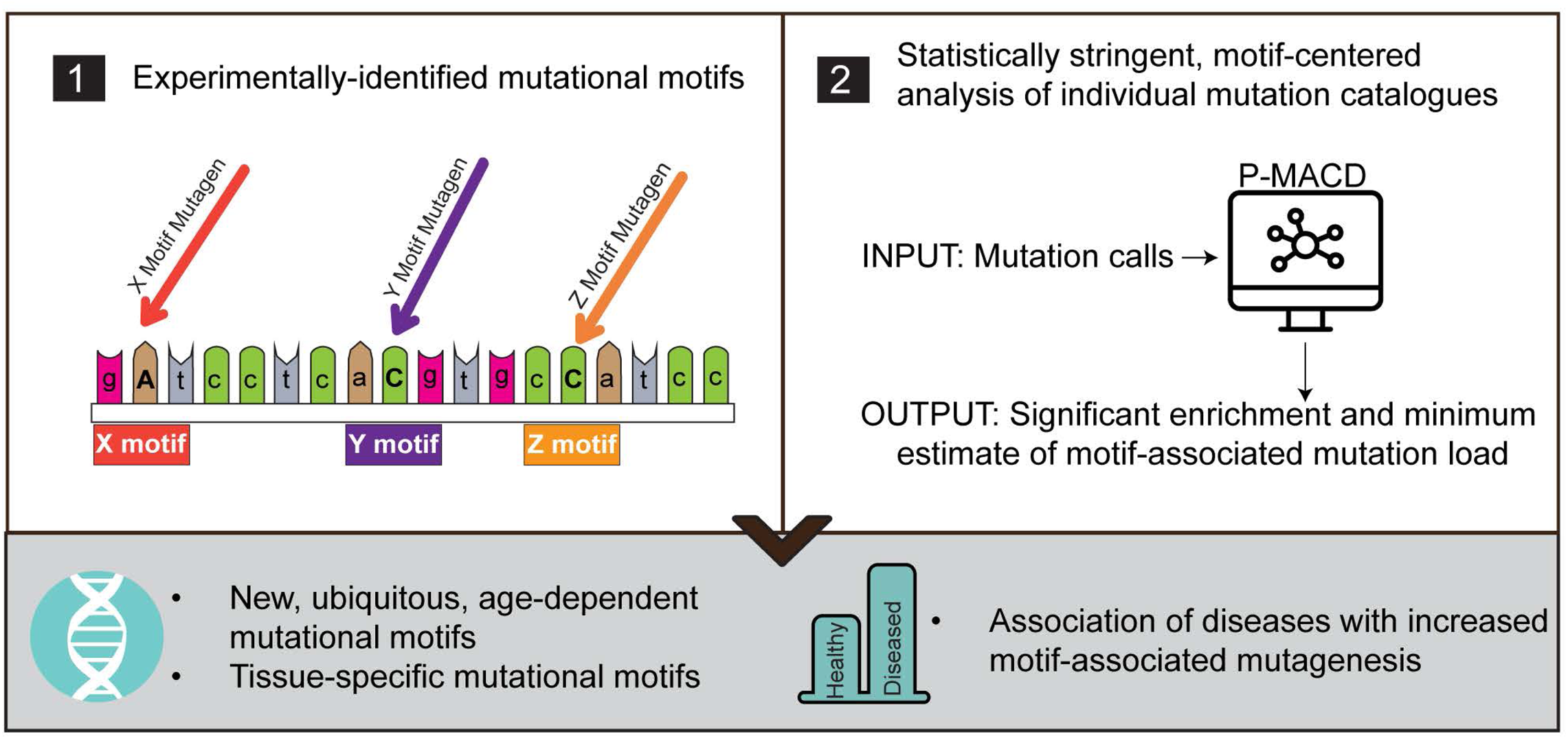

## References

1. Lawrence MS, Stojanov P, Polak P, et al. Mutational heterogeneity in cancer and the search for new cancer-associated genes. Nature. 2013;499(7457):214–8.

2. Roberts SA, Gordenin DA. Hypermutation in human cancer genomes: footprints and mechanisms. Nat Rev Cancer. 2014;14(12):786–800.

3. Alexandrov LB, Nik-Zainal S, Wedge DC, et al. Signatures of mutational processes in human cancer. Nature. 2013;500(7463):415–21.

4. Ames BN, Durston WE, Yamasaki E, Lee FD. Carcinogens are mutagens: a simple test system combining liver homogenates for activation and bacteria for detection. Proceedings of the National Academy of Sciences of the United States of America. 1973;70(8):2281–5.

5. Loeb LA, Springgate CF, Battula N. Errors in DNA replication as a basis of malignant changes. Cancer research. 1974;34(9):2311–21.

6. Erickson RP. Somatic gene mutation and human disease other than cancer. Mutation research. 2003;543(2):125–36.

7. Erickson RP. Recent advances in the study of somatic mosaicism and diseases other than cancer. Curr Opin Genet Dev. 2014;26:73–8.

8. Erickson RP. The importance of de novo mutations for pediatric neurological disease--It is not all in utero or birth trauma. Mutation research Reviews in mutation research. 2016;767:42–58.

9. Saini N, Gordenin DA. Somatic mutation load and spectra: A record of DNA damage and repair in healthy human cells. Environ Mol Mutagen. 2018;59(8):672–86.

10. Martincorena I, Campbell PJ. Somatic mutation in cancer and normal cells. Science. 2015;349(6255):1483–9.

11. Maslov AY, Vijg J. Somatic mutation burden in relation to aging and functional life span: implications for cellular reprogramming and rejuvenation. Curr Opin Genet Dev. 2023;83:102132.

12. Zhang L, Lee M, Maslov AY, Montagna C, Vijg J, Dong X. Analyzing somatic mutations by single-cell whole-genome sequencing. Nature protocols. 2024;19(2):487–516.

13. Evrony GD, Hinch AG, Luo C. Applications of Single-Cell DNA Sequencing. Annu Rev Genomics Hum Genet. 2021;22:171–97.

14. Shea A, Sun S, Kennedy J, Zhang L, Vijg J, Dong X. SomaMutDB 2.0: A comprehensive database for exploring somatic mutations and their functional impact in normal human tissues. Nucleic Acids Res. 2026;54(D1):D1291–d300.

15. Sun S, Wang Y, Maslov AY, Dong X, Vijg J. SomaMutDB: a database of somatic mutations in normal human tissues. Nucleic Acids Res. 2022;50(D1):D1100–d8.

16. Yaacov A, Rosenberg S, Simon I. Mutational signatures association with replication timing in normal cells reveals similarities and differences with matched cancer tissues. Scientific reports. 2023;13(1):7833.

17. Moore L, Cagan A, Coorens THH, et al. The mutational landscape of human somatic and germline cells. Nature. 2021;597(7876):381–6.

18. Alexandrov LB, Kim J, Haradhvala NJ, et al. The repertoire of mutational signatures in human cancer. Nature. 2020;578(7793):94–101.

19. Alexandrov LB, Nik-Zainal S, Wedge DC, Campbell PJ, Stratton MR. Deciphering signatures of mutational processes operative in human cancer. Cell reports. 2013;3(1):246–59.

20. Zhang T, Joubert P, Ansari-Pour N, et al. Genomic and evolutionary classification of lung cancer in never smokers. Nat Genet. 2021;53(9):1348–59.

21. Sui Y, Qi L, Zhang K, et al. Analysis of APOBEC-induced mutations in yeast strains with low levels of replicative DNA polymerases. Proceedings of the National Academy of Sciences of the United States of America. 2020;117(17):9440–50.

22. Gerhauser C, Favero F, Risch T, et al. Molecular Evolution of Early-Onset Prostate Cancer Identifies Molecular Risk Markers and Clinical Trajectories. Cancer Cell. 2018;34(6):996–1011 e8.

23. Ponomarev GV, Fatykhov B, Nazarov VA, et al. APOBEC mutagenesis is low in most types of non-B DNA structures. iScience. 2022;25(7):104535.

24. Kazanov MD, Roberts SA, Polak P, et al. APOBEC-Induced Cancer Mutations Are Uniquely Enriched in Early-Replicating, Gene-Dense, and Active Chromatin Regions. Cell reports. 2015;13(6):1103–9.

25. Henderson S, Chakravarthy A, Su X, Boshoff C, Fenton TR. APOBEC-mediated cytosine deamination links PIK3CA helical domain mutations to human papillomavirus-driven tumor development. Cell reports. 2014;7(6):1833–41.

26. Middlebrooks CD, Banday AR, Matsuda K, et al. Association of germline variants in the APOBEC3 region with cancer risk and enrichment with APOBEC-signature mutations in tumors. Nat Genet. 2016;48(11):1330–8.

27. de Bruin EC, McGranahan N, Mitter R, et al. Spatial and temporal diversity in genomic instability processes defines lung cancer evolution. Science. 2014;346(6206):251–6.

28. Zhang J, Fujimoto J, Zhang J, et al. Intratumor heterogeneity in localized lung adenocarcinomas delineated by multiregion sequencing. Science. 2014;346(6206):256–9.

29. Cortes-Ciriano I, Lee JJ, Xi R, et al. Comprehensive analysis of chromothripsis in 2,658 human cancers using whole-genome sequencing. Nat Genet. 2020;52(3):331–41.

30. Robertson AG, Kim J, Al-Ahmadie H, et al. Comprehensive Molecular Characterization of Muscle-Invasive Bladder Cancer. Cell. 2017;171(3):540–56 e25.

31. Hurst CD, Alder O, Platt FM, et al. Genomic Subtypes of Non-invasive Bladder Cancer with Distinct Metabolic Profile and Female Gender Bias in KDM6A Mutation Frequency. Cancer Cell. 2017;32(5):701–15 e7.

32. Consortium ITP-CAoWG. Pan-cancer analysis of whole genomes. Nature. 2020;578(7793):82–93.

33. Davis CF, Ricketts CJ, Wang M, et al. The somatic genomic landscape of chromophobe renal cell carcinoma. Cancer Cell. 2014;26(3):319–30.

34. Akdemir KC, Le VT, Kim JM, et al. Somatic mutation distributions in cancer genomes vary with three-dimensional chromatin structure. Nat Genet. 2020;52(11):1178–88.

35. Cancer Genome Atlas Research N, Albert Einstein College of M, Analytical Biological S, et al. Integrated genomic and molecular characterization of cervical cancer. Nature. 2017;543(7645):378–84.

36. Chan K, Roberts SA, Klimczak LJ, et al. An APOBEC3A hypermutation signature is distinguishable from the signature of background mutagenesis by APOBEC3B in human cancers. Nat Genet. 2015;47(9):1067–72.

37. Petljak M, Alexandrov LB, Brammeld JS, et al. Characterizing Mutational Signatures in Human Cancer Cell Lines Reveals Episodic APOBEC Mutagenesis. Cell. 2019;176(6):1282–94 e20.

38. Petljak M, Dananberg A, Chu K, et al. Mechanisms of APOBEC3 mutagenesis in human cancer cells. Nature. 2022;607(7920):799–807.

39. Zhang T, Sang J, Hoang PH, et al. APOBEC affects tumor evolution and age at onset of lung cancer in smokers. Nature communications. 2025;16(1):4711.

40. Saini N, Giacobone CK, Klimczak LJ, et al. UV-exposure, endogenous DNA damage, and DNA replication errors shape the spectra of genome changes in human skin. PLoS Genet. 2021;17(1):e1009302.

41. Hudson KM, Klimczak LJ, Sterling JF, et al. Glycidamide-induced hypermutation in yeast single-stranded DNA reveals a ubiquitous clock-like mutational motif in humans. Nucleic Acids Res. 2023;51(17):9075–100.

42. Lodato MA, Rodin RE, Bohrson CL, et al. Aging and neurodegeneration are associated with increased mutations in single human neurons. Science. 2017.

43. Miller MB, Huang AY, Kim J, et al. Somatic genomic changes in single Alzheimer’s disease neurons. Nature. 2022.

44. Lodato MA, Woodworth MB, Lee S, et al. Somatic mutation in single human neurons tracks developmental and transcriptional history. Science. 2015;350(6256):94–8.

45. Lee-Six H, Olafsson S, Ellis P, et al. The landscape of somatic mutation in normal colorectal epithelial cells. Nature. 2019;574(7779):532–7.

46. Ganz J, Luquette LJ, Bizzotto S, et al. Contrasting somatic mutation patterns in aging human neurons and oligodendrocytes. Cell. 2024;187(8):1955–70.e23.

47. Saini N, Roberts SA, Klimczak LJ, et al. The Impact of Environmental and Endogenous Damage on Somatic Mutation Load in Human Skin Fibroblasts. PLoS Genet. 2016;12(10):e1006385.

48. Blokzijl F, Janssen R, van Boxtel R, Cuppen E. MutationalPatterns: comprehensive genome-wide analysis of mutational processes. Genome Medicine. 2018;10(1):33.

49. Wang Y, Robinson PS, Coorens THH, et al. APOBEC mutagenesis is a common process in normal human small intestine. Nat Genet. 2023.

50. Li R, Du Y, Chen Z, et al. Macroscopic somatic clonal expansion in morphologically normal human urothelium. 2020;370(6512):82-9.

51. Díaz-Gay M, Vangara R, Barnes M, et al. Assigning mutational signatures to individual samples and individual somatic mutations with SigProfilerAssignment. Bioinformatics. 2023;39(12):btad756.

52. Roberts SA, Lawrence MS, Klimczak LJ, et al. An APOBEC cytidine deaminase mutagenesis pattern is widespread in human cancers. Nat Genet. 2013;45(9):970–6.

53. Wickham H. Data analysis. ggplot2: elegant graphics for data analysis: Springer; 2016. p. 189–201.

54. Hoppe J. Adobe Illustrator CC: a complete course and compendium of features. 1st edition ed. San Rafael, CA: Rocky Nook, Inc.; 2020.

55. You Y-H, Lee D-H, Yoon J-H, Nakajima S, Yasui A, Pfeifer GP. Cyclobutane Pyrimidine Dimers Are Responsible for the Vast Majority of Mutations Induced by UVB Irradiation in Mammalian Cells*. J Biol Chem. 2001;276(48):44688–94.

56. Pfeifer GP. Mutagenesis at methylated CpG sequences. Curr Top Microbiol Immunol. 2006;301:259–81.

57. Degtyareva NP, Placentra VC, Gabel SA, et al. Changes in metabolic landscapes shape divergent but distinct mutational signatures and cytotoxic consequences of redox stress. Nucleic Acids Res. 2023;51(10):5056–72.

58. Saini N, Sterling JF, Sakofsky CJ, et al. Mutation signatures specific to DNA alkylating agents in yeast and cancers. Nucleic Acids Res. 2020;48(7):3692–707.

59. Vijayraghavan S, Porcher L, Mieczkowski PA, Saini N. Acetaldehyde makes a distinct mutation signature in single-stranded DNA. Nucleic Acids Res. 2022.

60. Rosenquist TA, Grollman AP. Mutational signature of aristolochic acid: Clue to the recognition of a global disease. DNA Repair (Amst). 2016;44:205–11.

61. Li R, Di L, Li J, et al. A body map of somatic mutagenesis in morphologically normal human tissues. Nature. 2021;597(7876):398–403.

62. Olafsson S, McIntyre RE, Coorens T, et al. Somatic Evolution in Non-neoplastic IBD-Affected Colon. Cell. 2020;182(3):672–84.e11.

63. Roberts SA, Sterling J, Thompson C, et al. Clustered mutations in yeast and in human cancers can arise from damaged long single-strand DNA regions. Mol Cell. 2012;46(4):424–35.

64. Mertz TM, Collins CD, Dennis M, Coxon M, Roberts SA. APOBEC-Induced Mutagenesis in Cancer. Annu Rev Genet. 2022.

65. Peng W, Shaw BR. Accelerated deamination of cytosine residues in UV-induced cyclobutane pyrimidine dimers leads to CC-->TT transitions. Biochemistry. 1996;35(31):10172–81.

66. Nik-Zainal S, Alexandrov LB, Wedge DC, et al. Mutational processes molding the genomes of 21 breast cancers. Cell. 2012;149(5):979–93.

67. Saini N, Gordenin DA. Hypermutation in single-stranded DNA. DNA Repair (Amst). 2020;91–92:102868.

68. Dennen MS, Kockler ZW, Roberts SA, Burkholder AB, Klimczak LJ, Gordenin DA. Hypomorphic mutation in the large subunit of replication protein A affects mutagenesis by human APOBEC cytidine deaminases in yeast. G3 Genes|Genomes|Genetics. 2024;14(10):jkae196.

69. Sakofsky CJ, Saini N, Klimczak LJ, et al. Repair of multiple simultaneous double-strand breaks causes bursts of genome-wide clustered hypermutation. PLoS biology. 2019;17(9):e3000464.

70. Chan K, Gordenin DA. Clusters of Multiple Mutations: Incidence and Molecular Mechanisms. Annu Rev Genet. 2015;49:243–67.

71. Abascal F, Harvey LMR, Mitchell E, et al. Somatic mutation landscapes at single-molecule resolution. Nature. 2021;593(7859):405–10.

72. Brunner SF, Roberts ND, Wylie LA, et al. Somatic mutations and clonal dynamics in healthy and cirrhotic human liver. Nature. 2019;574(7779):538–42.

73. Yoshida K, Gowers KHC, Lee-Six H, et al. Tobacco smoking and somatic mutations in human bronchial epithelium. Nature. 2020.

74. Nishimura T, Kakiuchi N, Yoshida K, et al. Evolutionary histories of breast cancer and related clones. Nature. 2023;620(7974):607–14.

75. Chevalier A, Guo T, Gurevich NQ, Xu J, Yajima M, Campbell JD. Characterization of Mutational Signatures in Tumors from a Large Chinese Population. Cancer Research Communications. 2025;5(8):1466–76.

76. Díaz-Gay M, Zhang T, Hoang PH, et al. The mutagenic forces shaping the genomes of lung cancer in never smokers. Nature. 2025;644(8075):133–44.

77. Zanger UM, Schwab M. Cytochrome P450 enzymes in drug metabolism: regulation of gene expression, enzyme activities, and impact of genetic variation. Pharmacology & therapeutics. 2013;138(1):103–41.

78. Chen C, Wang DW. Chapter Seven - Cytochrome P450-CYP2 Family-Epoxygenase Role in Inflammation and Cancer. In: Hardwick JP, editor. Advances in Pharmacology: Academic Press; 2015. p. 193-221.

79. Kay J, Thadhani E, Samson L, Engelward B. Inflammation-induced DNA damage, mutations and cancer. DNA Repair. 2019;83:102673.

80. Cao H, Jiang Y, Wang Y. Kinetics of deamination and Cu(II)/H 2 O 2 /Ascorbate-induced formation of 5-methylcytosine glycol at CpG sites in duplex DNA. Nucleic Acids Research. 2009;37(19):6635–43.

81. Ren P, Zheng C, Pang Y, et al. Single-cell analysis of the somatic mutational landscape in human chondrocytes during aging and in osteoarthritis. Nature Aging. 2025;5(12):2417–31.

82. Sauty SM, Shine J, Bostan H, et al. Comparative whole-genome analyses of articular chondrocytes and skin fibroblasts reveal distinct genome instability landscapes in mesenchymal cell types. bioRxiv. 2026.

83. Hodgson SV, Foulkes WD, Maher ER, Turnbull C. Inherited Susceptibility to Cancer: Past, Present and Future. Annals of Human Genetics. 2025;89(5):354–65.

84. Robinson PS, Thomas LE, Abascal F, et al. Inherited MUTYH mutations cause elevated somatic mutation rates and distinctive mutational signatures in normal human cells. Nature Communications. 2022;13(1):3949.

85. Robinson PS, Coorens THH, Palles C, et al. Increased somatic mutation burdens in normal human cells due to defective DNA polymerases. Nature Genetics. 2021;53(10):1434–42.

86. Flynn A, Waszak SM, Weischenfeldt J. Somatic CpG hypermutation is associated with mismatch repair deficiency in cancer. Molecular systems biology. 2024;20(9):1006–24-24.

87. Palles C, West HD, Chew E, et al. Germline MBD4 deficiency causes a multi-tumor predisposition syndrome. Am J Hum Genet. 2022;109(5):953–60.

88. Querido I, Pinto C, Arinto P, et al. A Novel Deleterious Variant and a Founder Effect in Four New Families of MBD4-Associated Neoplasia Syndrome Recruited Over a Period of 20 Years. Clinical genetics. 2025.

89. Nik-Zainal S, Wedge DC, Alexandrov LB, et al. Association of a germline copy number polymorphism of APOBEC3A and APOBEC3B with burden of putative APOBEC-dependent mutations in breast cancer. Nat Genet. 2014;46(5):487–91.

90. Gansmo LB, Romundstad P, Hveem K, et al. APOBEC3A/B deletion polymorphism and cancer risk. Carcinogenesis. 2018;39(2):118–24.

